# Synaptic Zn^2+^ potentiates the effects of cocaine on striatal dopamine neurotransmission and behavior

**DOI:** 10.1101/2020.08.29.273482

**Authors:** Juan L. Gomez, Jordi Bonaventura, Jacqueline Keighron, Kelsey M. Wright, Dondre L. Marable, Lionel A. Rodriguez, Sherry Lam, Meghan L. Carlton, Randall J. Ellis, Chloe Jordan, Guo-hua Bi, Oscar Solis, Marco Pignatelli, Michael J. Bannon, Zheng-Xiong Xi, Gianluigi Tanda, Michael Michaelides

## Abstract

Cocaine binds to the dopamine (DA) transporter (DAT) to regulate cocaine reward and seeking behavior. Zinc (Zn^2+^) also binds to the DAT, but the *in vivo* relevance of this interaction is unknown. We found that Zn^2+^ concentrations in postmortem brain (caudate) tissue from humans who died of cocaine overdose were significantly lower than in control subjects. Moreover, the level of striatal Zn^2+^ content in these subjects negatively correlated with plasma levels of benzoylecgonine, a cocaine metabolite indicative of recent use. In mice, repeated cocaine exposure increased synaptic Zn^2+^ concentrations in the caudate putamen (CPu) and nucleus accumbens (NAc). Cocaine-induced increases in Zn^2+^ were dependent on the Zn^2+^ transporter 3 (ZnT3), a neuronal Zn^2+^ transporter localized to synaptic vesicle membranes, as ZnT3 knockout (KO) mice were insensitive to cocaine-induced increases in striatal Zn^2+^. ZnT3 KO mice showed significantly lower electrically-evoked DA release and greater DA clearance when exposed to cocaine compared to controls. ZnT3 KO mice also displayed significant reductions in cocaine locomotor sensitization, conditioned place preference (CPP), self-administration, and reinstatement compared to control mice and were insensitive to cocaine-induced increases in striatal DAT binding. Finally, dietary Zn^2+^ deficiency in mice resulted in decreased striatal Zn^2+^ content, cocaine locomotor sensitization, CPP, and striatal DAT binding. These results indicate that cocaine increases synaptic Zn^2+^ release and turnover/metabolism in the striatum, and that synaptically-released Zn^2+^ potentiates the effects of cocaine on striatal DA neurotransmission and behavior and is required for cocaine-primed reinstatement. In sum, these findings reveal new insights into cocaine’s pharmacological mechanism of action and suggest that Zn^2+^ may serve as an environmentally-derived regulator of DA neurotransmission, cocaine pharmacodynamics, and vulnerability to cocaine use disorders.

## Introduction

Zinc (Zn^2+^) is an essential trace element necessary for normal brain function (1–6). Within the brain it exists in two forms; a “fixed”, protein-bound form, which serves as a catalytic co-factor or as a structural component to Zn^2+^ binding proteins, and comprises ~90% of total brain concentration, or a “free”, or labile/chelatable form, comprising ~10% of total brain concentration. Labile Zn^2+^ is also referred to as “synaptic” Zn^2+^ as it is localized to synaptic vesicles in a subset of glutamatergic neurons, also referred to as “zincergic” neurons (7). Synaptic Zn^2+^ is found in several regions such as cortex (layers 2/3, 5 and 6), hippocampus, striatum, amygdala and is generally not present in thalamus (7, 8).

Synaptic Zn^2+^ levels are dependent on the Zn^2+^ transporter 3 (ZnT3) (9, 10), a neuronal-specific Zn^2+^ transporter localized to the membrane of synaptic vesicles. ZnT3 knockout (KO) mice lack the ability to package Zn^2+^ into these vesicles and consequently lack synaptic Zn^2+^ release (9, 10). ZnT3 KO mice are viable, fertile, and do not show major behavioral abnormalities across spatial learning, memory, or sensorimotor tasks, though they do exhibit small deficits in skilled reaching tasks (11, 12), fear learning (13), and sensory deficits (14, 15).

In terms of function, synaptic Zn^2+^ is released upon neuronal activation, serves as a key regulator of neurotransmitter signaling, and as such has also been referred to as a neurotransmitter (1–6). Tonic extracellular Zn^2+^ concentrations are estimated to be at <25 nM (16). However, upon physiological zincergic neuron activation and subsequent release, phasic Zn^2+^ concentrations can increase up to 10 μM (17). In this way, Zn^2+^ is thought to exert various effects on neurotransmission by binding to synaptic proteins that contain either low- or high-affinity Zn^2+^ binding sites (6). One such protein is the dopamine transporter (DAT), which contains four high-affinity Zn^2+^ binding sites on its extracellular domain (18–22).

Studies have shown that cocaine binds to the DAT to inhibit synaptic DA reuptake, which leads to an increase of extracellular DA (23). This mechanism underlies the direct subjective responses that accompany cocaine use (24), and is critical to cocaine self-administration in laboratory models, and cocaine reward and abuse liability in humans (25). *In vitro* assays have shown that when Zn^2+^ binds to the DAT it promotes a conformation that inhibits DA uptake and, when cocaine is present, Zn^2+^ can increase cocaine’s affinity and can modulate its potency to inhibit DA uptake (18–22). Interestingly, synaptic Zn^2+^ is present in the striatum (26, 27) where the DAT is found in highest concentration (28), and synaptic Zn^2+^ depletion in this region improves motor deficits and memory impairments caused by loss of striatal DAergic fibers (29). These prior studies suggest that synaptic Zn^2+^ may modulate *in vivo* DAT function and cocaine-DAT pharmacodynamic interactions in the striatum. However, the extent to which such interactions occur is unclear.

In addition to ZnT3, synaptic Zn^2+^ levels also depend on the environmental availability of Zn^2+^ which is obtained exclusively via the diet. Consequently, dietary factors which limit Zn^2+^ intake influence both synaptic Zn^2+^ levels and ZnT3 expression (30, 31). In fact, as the body lacks a specialized system for its storage, Zn^2+^ needs to be consumed continuously to avoid a state of deficiency which itself can have profound changes in both the periphery as well as in brain function (32). Human drug users generally do not follow a life-style that prioritizes nutritional needs and are known to exhibit nutritional deficiencies including dysregulated blood and hair Zn^2+^ content (33–38). However, whether such deficits are involved in the neurobiological changes associated with cocaine use or use of other drugs is unknown.

## Methods

### Subjects

De-identified postmortem human brain specimens were collected during the routine autopsy process as described in detail previously (39, 40). Briefly, the cause and manner of death were determined by forensic pathologists following medico-legal investigations that evaluated the circumstances of death including medical records, police reports, autopsy results, and toxicological data. Inclusion in the cocaine cohort (n = 20; 10 Caucasian, 10 African-American) was based on cocaine abuse as the cause of death, a documented history of drug abuse, and a toxicology positive for high levels of the cocaine metabolite benzoylecgonine and, in most cases, the short-lived cocaine adulterant levamisole, both indicative of recent cocaine use prior to death. Control subjects (n=20; 10 Caucasian, 10 African-American) died as a result of cardiovascular disease or gunshot wound, had no documented history of drug abuse, and tested negative for cocaine and other drugs of abuse. Exclusion criteria for the study included a known history of neurological or psychiatric illness, death by suicide, estimated postmortem interval exceeding 20 hr, evidence of neuropathology (e.g. encephalitis, stroke), or chronic illness (e.g. cirrhosis, cancer, HIV, prolonged hospitalization). The final groups did not differ with regard to age for race, nor with regard to brain pH, a well-established measure of sample quality and perimortem agonal state (41). Tissue from one cocaine user was not included due to very high cocaine metabolite levels.

Given that the human tissue samples available consisted only of males, and therefore all human data were limited to this sex, we focused on male mice for all other mechanistic experiments. Male C57Bl/6J mice were acquired from Jackson Labs at 8-weeks of age. Breeding pairs of *Slc30a3* (ZnT3) knockout mice were obtained from Dr. Thanos Tzounopoulos at the University of Pittsburgh and bred at the National Institute on Drug Abuse (NIDA) (Baltimore, MD) on a C57Bl/6J background. Mice were genotyped by Transnetyx (Cordova, TN). All mice were male and matched for age and weight. Mice were single-housed during experimental testing in a temperature and humidity-controlled environment on a regular light cycle (on at 7 am and off at 7 pm). Food and water were available ad libitum and mice were acclimated prior to any behavioral procedures by handling. All experimental procedures were carried out in accordance with the National Institutes of Health Guide for the Care and Use of Laboratory Animals and were approved by the Animal Care and Use Committee of NIDA.

### Total Reflection X-ray Spectroscopy (TXRF)

Tissue samples were collected and weighed in 1.5 mL Eppendorf tubes. The weight of the tissue was directly used to calculate element concentrations in μg/kg units. Each tissue sample was dissolved in 100 μL of nitric acid (Sigma: NX0408) with 2 μL of a gallium standard (conc. 1000 ppm). Each sample was assessed in duplicate for TXRF elemental analysis using an S2 Picofox (Bruker, Billerica, MA). This instrument exposes the sample to an X-ray beam and measures fluorescence radiation specific to the element(s) of interest. Human samples were prepared from postmortem tissue collected from the anterior caudate. Wild-type (WT) or ZnT3 KO mice received either saline, a single cocaine injection (20 mg/kg, i.p), or 8 repeated daily cocaine (20 mg/kg, i.p) injections and were killed 24 hrs after the last injection. Mice fed the 30 ppm and 5 ppm Zn^2+^ diets were exposed to each respective diet for 35 days and then killed. Mouse samples were prepared by slicing flash frozen tissue on a cryostat (100 μm sections from Bregma 1.00 mm to 0.00 mm) or dissecting the cortex and striatum from each section.

### Synchrotron X-ray Fluorescence Microspectroscopy (μXRFS)

Brain concentrations and distributions of Zn^2+^ from C57BL/6J mice injected with saline or cocaine (10 mg/kg, i.p.) every other day for 8 days and killed 24 hrs after the last injection were measured at the X26a beamline at the National Synchrotron Light Source (NSLS) at Brookhaven National Laboratory (Upton, NY). The synchrotron X-ray beam was tuned to 12 keV using a Si(111) channel-cut monochromotor. The monochromatic beam was then collimated to 350 μm×350 μm and then focused to approximately 6 μm×10 μm using Rh-coated silicon mirrors in a Kirkpatrick–Baez (KB) geometry. The sample was placed at a 45° angle to the incident X-ray beam and X-ray fluorescence was detected with an energy dispersive, 9-element germanium array detector (Canberra, Meriden, CT) oriented at 90° to the incident beam. The sample was approximately 6 cm from the detector. A light microscope objective (Mitutoyo, M Plan Apo 5X) was coupled to a digital CCD camera for sample viewing. Energy dispersive spectra were collected by raster-scanning the sample through the X-ray beam using a dwell time of 0.3 s/pixel and a step size of 10 μm. Zn Kα, fluorescence counts were then extracted from background-corrected energy dispersive spectra. All data were normalized to variations in incident photon flux by normalizing to changes in I0 measured by ion chamber upstream of the KB optics. XRFS calibration standards on Nuclepore® polycarbonate aerosol membranes expressing known (±5%) concentrations of Zn (48.4 μg/cm^2^) were also imaged in parallel to the samples (Micromatter, Vancouver, BC) and used to express results as μg/cm^2^. Image analysis was carried out using ImageJ (National Institutes of Health, Bethesda, MD). Regions of interest (ROI) were drawn onto the cortex (Ctx), caudate putamen (CPu), nucleus accumbens (NAc), and measurements for each ROI were obtained.

### Zinc-Selenium Autometallography (ZnSe^AMG^)

WT and ZnT3 KO mice were anesthetized with a ketamine-xylazine cocktail (ket=60mg/kg + xyl=12mg/kg) and injected (i.p.) with 15 mg/kg sodium selenite (Sigma Aldrich: 214485) and placed on a heating pad while anesthetized for 60 minutes. Mice were then perfused with 0.1M phosphate buffer for 5 minutes. Brain tissue was dissected, and flash frozen in dry ice cooled isopentane and stored at −80°C until sectioning. Coronal brain sections (20 μm) were thaw mounted at the level of the striatum and hippocampus on positively-charged glass slides. Slides were stored at −20°C until staining. Slides were loaded in non-metallic staining racks and allowed to reach room temperature. Slides were fixed in 95% ethanol for 15 min followed by hydration in 70% (2 min) and 50% (2 min) ethanol ending in 3 × 2 min distilled water rinses. Slides were dipped in 0.5% gelatin and air dried prior to physical development. Developer was made by mixing Gum Arabic (50% solution, 100mL), citrate buffer (2.0M, 20mL), hydroquinone (1.7g in 30mL DDH_2_O), silver lactate (0.22g in 30mL H_2_0), and DDH_2_O (200mL). Developer was poured onto slides, incubated for 60 min in the dark then quickly checked at 10 min intervals until sections are dark brown. Slides were washed in slowly flowing tap water (37°C) for 10 min to remove gelatin then rinsed 3 × 2 min in distilled water. Slides were then incubated in 5% sodium thiosulphate (12min) and rinsed 2 × 2min in distilled water and post-fixed in 70% ethanol (30min). Optional counter stain using cresyl violet or toluidine blue (5min) followed by rinse 4 × 30 sec rinse in distilled water. Finally, slides were dehydrated in 95% ethanol (5min), 100% ethanol 2 × 5 min, xylene 2 × 5 min, and coverslipped with permount. Stained sections were imaged using brightfield microscopy.

### ^65^Zn Uptake Experiments using Positron Emission Tomography (PET)

WT and ZnT3 KO mice were anesthetized with isoflurane and placed in a prone position on the scanner bed of a nanoScan PET/CT (Mediso, USA) injected intravenously (~150 μL) with ^65^ZnCl_2_ (~2.2 MBq) and PET data were acquired for 2 hours followed by a CT scan. After scanning, animals were returned to their home cage. Scans were repeated on days 1, 3, 7, and 14. For cocaine experiments, C57Bl/6J mice were injected with ^65^ZnCl_2_ as above and then injected immediately with saline or cocaine (20 mg/kg, i.p). Saline and cocaine injections continued daily for 7 days. Mice were scanned on Day 1 and Day 7 after ^65^ZnCl_2_ injection as above. In all cases, the PET data were reconstructed and corrected for dead-time and radioactive decay. Qualitative and quantitative assessments of PET images were performed using the PMOD software environment (PMOD Technologies, Zurich Switzerland). Time-activity curves were generated using manually drawn volumes of interest using the CT image as a reference. Standardized uptake values (SUV) were calculated using the formula SUV(i) = C(i) / (ID × BW) where C(i) is the activity value at a given time point (in kBq/cc), ID is the injected dose (in MBq) and BW is the animal’s body weight (in kg). For voxel-wise analyses we used Statistical Parametric Mapping (SPM12, London, UK) as previously described (42). First, all the images were co-registered and masked to the reference mouse atlas in PMOD. Regional changes in uptake were assessed relative to global (whole-brain) uptake. For the cocaine experiments, a repeated measures analysis of variance (ANOVA) model was used that defined saline vs. cocaine-treated mice scanned at 1- and 7-days post ^65^ZnCl_2_ injection. Images were subtracted after intensity normalization to 100 by the proportional scaling method. After estimation of the statistical model, a contrast (Cocaine > Vehicle) was applied to reveal the effects of interest. These effects were overlaid on the reference MRI. An uncorrected *P*-value of 0.05 with a cluster threshold value of 50 were used as thresholds to determine statistical significance.

### *Ex vivo* ^65^Zn autoradiography

One day after the last PET scan, WT and ZnT3 KO mice were killed and brain tissue was dissected, flash frozen in isopentane, and stored at −80°C until sectioning. Tissue was sectioned and thaw mounted on positively charged glass slides. Slides were placed on BAS-IP SR 2040 E Super Resolution phosphor screens (GE Healthcare) for 14-days and imaged using a phosphor imager (Typhoon FLA 7000; GE Healthcare).

### Radioligand Binding Assays

Brains from killed C57Bl/6J mice were removed and striata dissected and quickly frozen until use. The tissue was weighed and suspended in 10 times (w/v) of ice-cold Tris-HCl buffer (50 mM, pH 7.4). The suspension was homogenized with a Polytron homogenizer (Kinematica, Basel, Switzerland) under ice. Homogenates were centrifuged at 48,000g (50 min, 4 °C) and washed twice in the same conditions to isolate the membrane fraction. Protein was quantified by the bicinchoninic acid method (Pierce). For competition experiments, membrane suspensions (50 μg of protein/ml) were incubated in 50 mM Tris-HCl (pH 7.4) 0.5 nM of [^3H^]WIN-35428 (Perkin-Elmer) and increasing concentrations of the indicated competing drugs (WIN-35428 or cocaine) in the presence or the absence of 100 nM, 10 μM or 1 mM of ZnCl_2_ during 2 h at RT. Nonspecific binding was determined in the presence of 100 μM cocaine. In all cases, free and membrane-bound radioligand were separated by rapid filtration through Whatman (Clifton, NJ) GF/B filters, pre-soaked in 0.05% polyethyleneimine by using a Brandel R48 filtering manifold (Brandel Inc., Gaithersburg, MD). The filters were washed twice with 5 ml of cold buffer and transferred to scintillation vials. Beckman Ready Safe scintillation cocktail (3.0 ml) was added, and the vials were counted the next day with a Beckman 6000 liquid scintillation counter (Beckman Coulter Instruments, Fullerton, CA) at 50% efficiency.

### *In vitro* Autoradiography using [^3^H]WIN-35,428

Brain tissue from WT and ZnT3 KO mice was dissected, flash frozen in isopentane, and stored at −80°C until sectioning. Tissue was sliced on a cryostat at 16 μm and thaw mounted on positively charged glass slides and stored at −20°C until autoradiography. Incubation Buffer consisted of 50 mM Tris-HCl (7.4 pH) and 100 mM NaCl in deionized water. [^3^H]WIN-35,428 Total binding buffer (S.A. 82.9 Ci/mmol, Conc. 1 mCi/mL) was made in incubation buffer at a concentration of 10 nM. ZnCl_2_ binding buffer was made using the Total binding buffer stock and adding ZnCl_2_ for a concentration of 10 μM. Slides were pre-incubated in ice-cold incubation buffer for 20 minutes then transferred to respective radioactive incubation buffers (i.e. total or total+ZnCl_2_) for 120 minutes on ice. Slides were then washed 2 × 1min in ice-cold 50 mM Tris-HCl (pH=7.4) then dipped (30 sec) in ice-cold deionized water. Slides were dried under stream of cool air and placed on BAS-IP TR 2025 E Tritium Screen (GE Healthcare) for 5-days and imaged using a phosphor imager (Typhoon FLA 7000; GE Healthcare). Sections were analyzed using Multigauge software (Fujifilm, Japan).

### *In vivo* Fast Scan Cyclic Voltammetry (FSCV)

FSCV procedures follow those of recently published work from our laboratory in anesthetized mice using electrical stimulation (43). Briefly, glass sealed 100 μm carbon fiber microelectrodes were pre-calibrated with known concentrations of dopamine and changes in pH to allow for a principal component analysis (PCA) of the raw data using HDCV (UNC, Chapel Hill, NC). Dopamine was identified by cyclic voltammogram using a voltage scan from −0.3 to 1.4V at 400 V/s. During the experiment an external stimulus was applied using the tungsten electrode every 5 min comprised of 24 pulses 4ms in width at 60 Hz and 180 μA while the working electrode was implanted in the striatum (AP: +1.5 mm; ML: ±1.0 mm; DV: −3.2 to −3.7 mm from bregma). After PCA data were analyzed to determine the DA_Max_ and DA clearance rate using a custom macro written in Igor Carbon Pro which identified peaks greater than 3x root mean square noise and fit to equation 1 where DA_Max_ represents the peak DA concentration measured, k is the rate constant, and t is time (43).

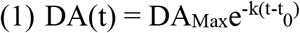

### Cocaine Locomotor Sensitization

Each session during the development phase was 30 minutes and mice were only exposed to one session per day with locomotor activity quantified as distance traveled (cm). All injections were administered (i.p.). Mice were first habituated to the locomotor activity chambers (Opto-varimex ATM3, Columbus Instruments). On the next two sessions mice were injected with saline and placed in the chambers. On the following five sessions, separate groups were injected with either saline or cocaine (10 mg/kg) in a counterbalanced design. Mice were then allowed 7-days of withdrawal in the colony room and then returned to the behavior room for testing expression of sensitization. Briefly, all mice were allowed access to the activity chambers for 60 minutes followed by increasing doses of cocaine (saline, 5, 10, 20 mg/kg) every 60 minutes. Data collection was paused but chambers were not cleaned in-between cocaine dosing, each mouse was picked up, injected, and placed back in chamber to continue data collection.

### Cocaine Conditioned Place Preference (CPP)

The task consisted of 10 sessions, 1 per day, in chambers with two visually distinct sides, one with clear walls and white floor and one with checkered walls and black floor. The sides were divided by a door with and without access to the other side. Locomotor activity was measured by way of time spent in each chamber as well as total distance traveled (Opto-varimex ATM3, Columbus Instruments). In the first session the mice could explore both sides of a conditioning box for 15 minutes to determine inherent side preference, designated as the Pre-Test. Using this data, the cocaine-paired side was pseudo-randomized so that mice with a preference for one side (>60%) were cocaine-paired on the other, non-preferred side. Mice with no side preference were cocaine-paired in a counterbalanced fashion. Separate groups of mice were conditioned with either a 5, 10, or 20 mg/kg dose of cocaine. In an alternating fashion for 8-days, mice were injected (i.p.) with either saline or cocaine and placed in the predetermined drug/no drug side of the chamber for 30 minutes. The mice had no physical access to the other side but were still able to see through the clear divider wall. Each mouse had a total of 4 saline-paired days and 4 cocaine-paired days. The last session was the same as the first and designated the Test session. Time spent in the cocaine-paired chamber during the Pre-Test session (prior to conditioning) was subtracted from time spent in the cocaine-paired chamber during the Test session and expressed as the Preference score.

### Mouse Intravenous Cocaine Self-Administration

Mice were implanted with jugular vein catheters under ketamine/xylazine anesthesia and using aseptic surgical techniques. A 6.0 cm length MicroRenathane (ID 0.012”, OD 0.025”; Braintree Scientific Inc., Braintree, MA, USA) catheter was inserted 1.2 cm into the right jugular vein and anchored to a 24-gauge steel cannula (Plastics One, Roanoke, VA, USA) that was bent at a 100° angle and mounted to the skull with cyanoacrylate glue and dental acrylic. A 2.5-cm extension of flexible tubing was connected to the distal end of the cannula. The mice were allowed 5–7 days for recovery, during which time 0.05 ml of a 0.9% saline solution containing 20 IU/ml heparin and 0.33 mg/ml gentamycin was infused daily through the catheter to prevent catheter clotting and infection. Thereafter, 0.05 ml of 0.9% saline solution containing 20 IU/ml heparin was infused immediately prior to and immediately following each daily self-administration session. When needed, i.v. brevital (a barbiturate) was used to test catheter patency between the self-administration sessions. During cocaine self-administration sessions, the flexible tubing extension was connected to a perfusion pump (Med Associates, Fairfax, VT) via a PE50 tubing connector. After daily self-administration sessions, the free end of the cannula guide was always kept sealed.

Operant test chambers (Med Associates, Fairfax, VT) contained two levers (active and inactive) located 2.5 cm above the floor as well as a cue light above each lever. A house light mounted on the opposite side of the chamber signaled the start of each 3 hr session and remained illuminated until the session ended. For self-administration sessions, a liquid swivel mounted on a balance arm above the chamber allowed for i.v. drug delivery in freely-moving mice. Depression of the active lever resulted in the activation of an infusion pump; depression of the inactive lever was recorded but had no scheduled consequences. Each infusion was paired with two discrete cues: illumination of the cue light above the active lever, and a cue tone that lasted for the duration of the infusion. Experimental events were controlled by a PC programmed in Medstate Notation and connected to a Med Associates interface.

After recovery from surgery, mice were placed into operant chambers and allowed to lever-press for i.v. cocaine self-administration under a fixed-ratio 1 (FR1) reinforcement schedule (i.e., each lever press leads to one cocaine infusion) for 3 h daily. Each cocaine infusion lasted 4.2 sec, during which additional active lever responses were recorded but had no consequences (i.e., non-reinforced active lever response). Mice were trained initially for a high unit dose of cocaine (1 mg/kg/infusion) to potentiate acquisition of self-administration until stable self-administration was achieved, which was defined as earning at least 20 infusions per 3 hr session and an active/inactive lever press ratio exceeding 2:1. Then the mice were switched to a multiple-dose schedule to observe the dose-dependent cocaine self-administration according to a descending cocaine dose sequence from the initial dose of 1 mg/kg/infusion (sessions 1-13) to 0.5 mg/kg/infusion (sessions 14-20), 0.25 mg/kg/infusion (sessions 21-23), 0.125 mg/kg/infusion (sessions 24-27), and 0.0625 mg/kg/infusion (sessions 28-29). Mice that did not reach stability criteria, lost catheter patency, or showed excessive high-level inactive lever responding (>100 presses per session) were excluded from further experimentation. To prevent cocaine overdose, maximally allowed cocaine infusions were 50 (0.1 and 0.5 mg/kg/infusion), 100 (0.25 mg/kg/infusion), 200 (0.125 mg/kg/infusion), or 400 (0.0625 mg/kg/infusion), respectively during each 3-h session. The number of cocaine infusions earned, and active and inactive lever responses were recorded for each session. The last 2-3 days of cocaine self-administration data at each dose were averaged and used to compare dose-response performance between WT and KO mice.

After the completion of the above cocaine dose-response experiment, the animals were switched to cocaine self-administration under PR reinforcement schedule. During PR conditions, the work requirement (lever presses) needed to receive a cocaine infusion was raised progressively within each test session according to the following PR series: 1, 2, 4, 6, 9, 12, 15, 20, 25, 32, 40, 50, 62, 77, 95, 118, 145, 178, 219, 268, 328, 402, 492, and 603 until the break point was reached. The break point was defined as the maximal workload (i.e., number of lever presses) completed for the last cocaine infusion prior to a 1-h period during which no infusions were obtained by the animal. Animals were tested for cocaine self-administration under PR reinforcement at three doses (starting at 0.25, then 1 and then 0.5 mg/kg/infusion) from days 30 to 38.

After the completion of the PR experiments, the same groups of animals continued for cocaine extinction and reinstatement tests. During extinction, syringe pumps were turned off and the cocaine-associated cue light and tone were unavailable. Thus, lever pressing was recorded but had no scheduled consequences. Extinction training continued for about 20 days until the extinction criteria were met (i.e., lever responding <20% of the self-administration baseline) for at least 3 sessions. Mice then received a 10 mg/kg i.p cocaine injection to evoke reinstatement of drug-seeking behavior. During reinstatement testing, active lever presses lead to re-exposure to the cue light and tone previously paired with cocaine infusions, but not to actual cocaine infusions. Active and inactive lever responses were recorded for each extinction and reinstatement session. Lever pressing behavior during the cocaine-primed session was compared to the average lever pressies during the last 3 days of extinction.

### Custom Diets

Diets were formulated by Research Diets, Inc via use of AIN-93M mature rodent diet. The diets were compositionally identical, but one diet had an adequate amount of Zn^2+^ (30 ppm) and the other diet had a deficient amount of Zn^2+^ (5 ppm). Zn^2+^ concentration in each diet was confirmed in-house via random sampling of chow pellets and TXRF (S2 Picofox, Bruker, Billerica, MA).

### Body Weight and Food Intake Measurements

C56BL/6J mice arrived at the NIDA mouse colony and were allowed one week of environmental acclimation with regular chow and water available *ad lib*. After the acclimation period, mice were given a diet containing either 30 ppm Zn^2+^ or 5 ppm Zn^2+^ for a minimum of 35 days prior to any behavioral manipulations. Mice stayed on the diet until the completion of the experiment. Mice were individually housed and weighed 3 times per week (MWF) along with food weight to track the amount of food consumed.

### Immunohistochemistry

Coronal sections were sliced on a cryostat (30 μm), collected in 6-well plates with PBS, and stored at 4°C until use. Sections were transferred to 12-well plates and permeabilized in washing buffer (PBS + Triton X-100 0.1%) for 10 min at room temperature on shaker. Tissue was blocked in blocking buffer (BSA 3% + PBS + Triton X-100 0.1%) for 60 min at RT on shaker. Tissue was incubated overnight in primary ZnT3 antibody (1:500) (anti-rabbit, #197 003, Synaptic Systems, Goettingen, Germany) at 4°C. Tissue was washed with washing buffer 3 × 10 min at RT then incubated in secondary antibody (Alexa 488-Rabbit #A1134, Thermo Fisher (1:400), Topro (1:1200), and DAPI (1:600) for 2 hrs at RT in the dark. Tissue was washed with washing buffer 3 × 10 min, transferred to dish with PBS, mounted on positively charged glass slides, and coverslipped with aqueous mounting medium (90% glycerol + 30 mM Tris-HCl, pH 8.0) and imaged using confocal microscopy.

### Statistics

Sample sizes were estimated based on prior experience with the specific assays/experiments and expected effect sizes. Mice (with the exception of ZnT3 KO) were randomly selected from littermates into experimental groups. Experiments were performed unblinded. Depending on experiment, we used linear regression, unpaired t-tests, one sample non-parametric (Wilcoxon) tests, one or two-way ANOVA or a mixed effects model (to account for missing data) taking repeated measures (RM) into account when appropriate. Significant main or interaction effects were followed by Holm-Sidak pairwise comparisons. All statistical tests were evaluated at the p≤0.05 level. Estimations statistics were used for experiments with low sample size (e.g. n<5) or with data that were not normally distributed (www.estimationstats.com)(44).

## Results

### Striatal Zn^2+^ is low in human cocaine users and correlates with cocaine intake

The highest DAT density in the brain is found in the striatum (28), a region heavily implicated in cocaine addiction (45–48). Human drug users are characterized by nutritional deficiencies and dysregulated blood and hair Zn^2+^ content (33–38) but whether they show deficits in striatal Zn^2+^ is unknown. To assess this, we performed elemental profiling using total reflection X-ray fluorescence spectroscopy (TXRF) in postmortem striatal tissue derived from humans whose primary cause of death was attributed to cocaine abuse (n=19) or matched controls (n=20) (Figure 1A). Cocaine users and controls did not differ in age (Figure 1B) or brain pH content (Figure 1C). We found that cocaine users had significantly lower (unpaired t-test, t=2.87; p=0.006) striatal Zn^2+^ levels compared to controls (Figure 1D). This difference was selective to Zn^2+^ as we did not detect significant differences in other elements (Figure S1). Striatal Zn^2+^ levels in these subjects significantly and negatively correlated (linear regression; F(1, 17)=4.5; p=0.04) with plasma concentrations of benzoylecgonine, (Figure 1E) a stable cocaine metabolite indicative of recent cocaine use (49).

**Figure 1.**
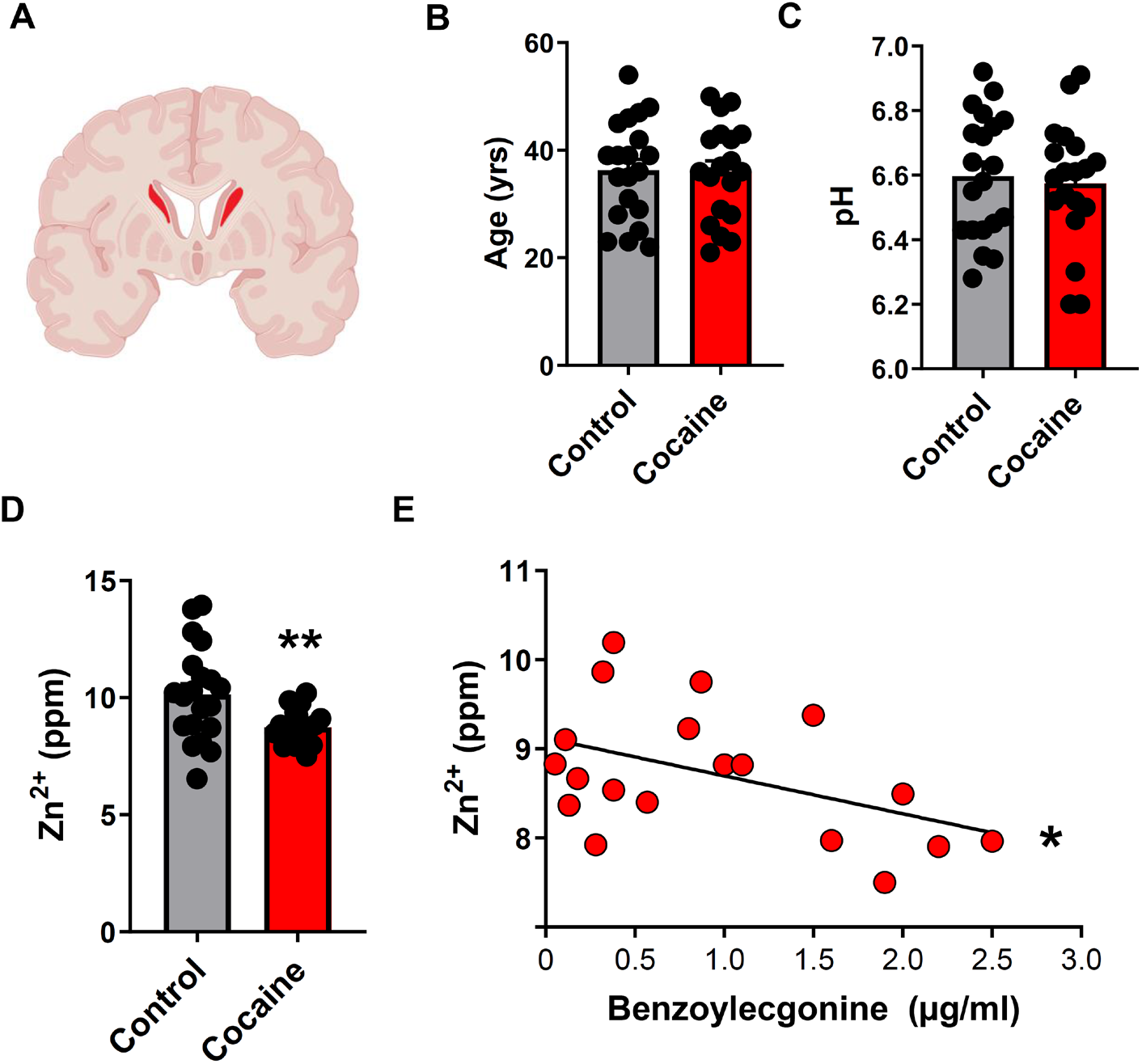
Striatal Zn^2+^ is low in human cocaine users and correlates with cocaine intake. (**A**) Schematic showing sampled region (caudate) from postmortem human brain samples. (**B**) Cocaine users and control subjects did not differ in age or (**C**) in tissue pH. (**D**) Cocaine users showed significantly lower striatal Zn^2+^ levels compared to control subjects. (**E**) Striatal Zn^2+^ levels in cocaine users significantly correlated with plasma benzoylecgonine levels (ppm – parts per million). **p≤0.01, *p≤0.05. All data expressed as Mean ±SEM.

### Cocaine increases synaptic Zn^2+^ release and uptake in the striatum

To obtain mechanistic insights in the relationship between cocaine exposure and striatal Zn^2+^ levels, we examined striatal Zn^2+^ content in mice exposed to cocaine. Unlike TXRF, synchrotron X-ray fluorescence microspectroscopy (μSXRF) allows visualization and quantification of Zn^2+^ in brain slices (50, 51). We first verified that the distribution of Zn^2+^, as detected by μSXRF in the hippocampus, where synaptic Zn^2+^ levels are high, coincided with distributions of ZnT3 immunolabeling and histochemically-reactive synaptic Zn^2+^ (52) (Figure S2). We then performed μSXRF in striatal sections from mice exposed to daily cocaine injections (10 mg/kg, intraperitoneal (IP), 4 days, n=4) and killed 24 hours later. We found that cocaine-exposed mice had significantly greater total Zn^2+^ levels in the caudate putamen (CPu) [unpaired mean difference: Veh vs. Coc=111; (95.0% CI 66.4, 139), p=0.003] and in the nucleus accumbens (NAc) [unpaired mean difference: Veh vs. Coc=65.3; (95.0% CI 47.4, 98.2); p<0.001] compared to vehicle-injected mice (n=5) (Figure 2, A and B). There was no significant difference in cortex [unpaired mean difference: Veh vs. Coc=77.8; (95.0% CI 85.1, 141); p=0.09].

**Figure 2.**
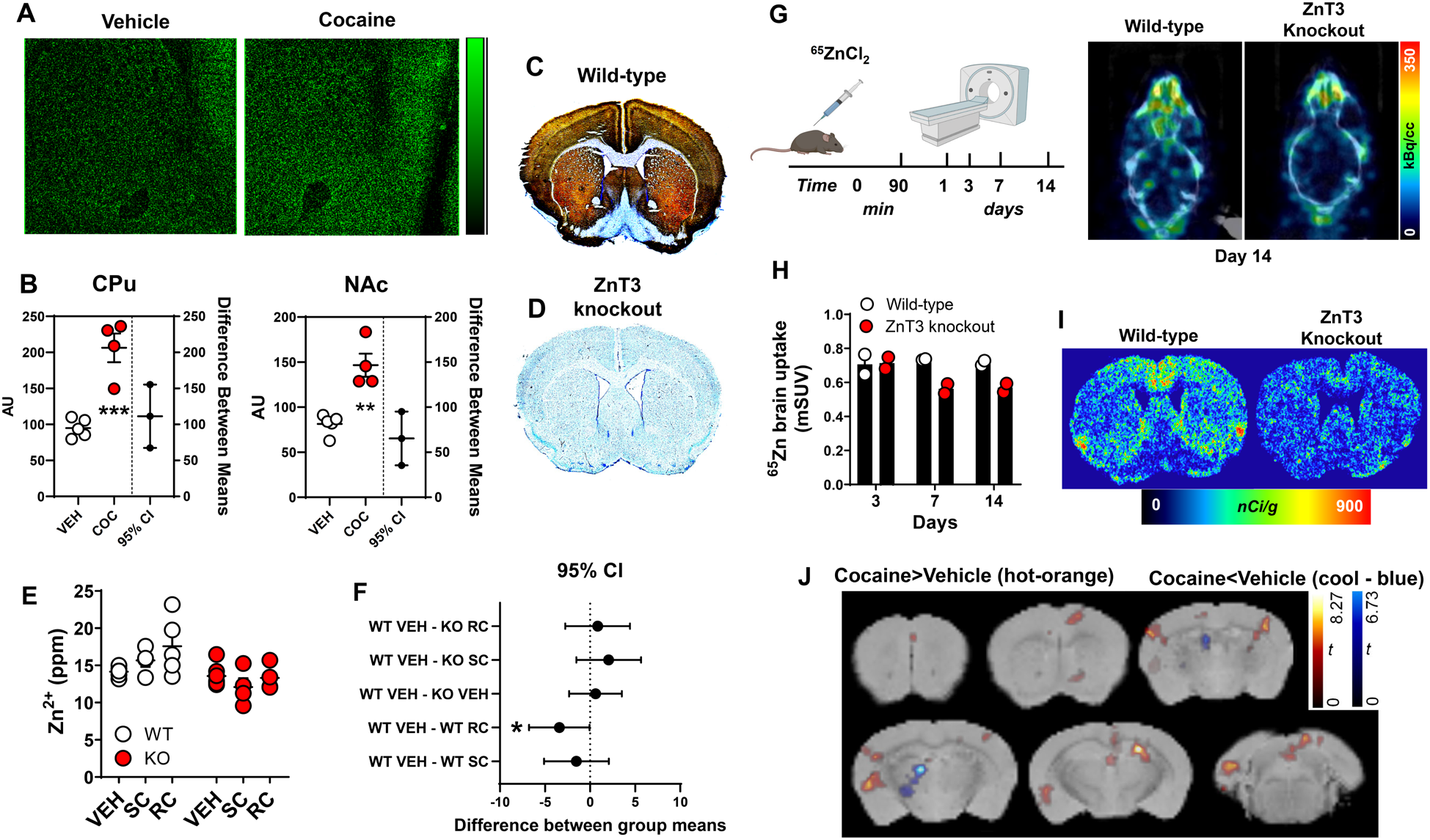
Cocaine increases synaptic Zn^2+^ release and uptake in the striatum. (**A**) Representative synchrotron X-ray fluorescence microspectroscopy (μSXRF) Zn^2+^ maps from vehicle- or cocaine-treated mice. (**B**) Cocaine-treated mice had significantly greater Zn^2+^ (AU-arbitrary units) than vehicle-treated mice in caudate putamen (CPu) and nucleus accumbens (NAc). (**C**) Representative Timm- and cresyl violet-co-stained sections from a wild-type (WT) and (**D**) a ZnT3 knockout (KO) mouse. (**E**) Zn^2+^ content measured using TXRF in wild-type and ZnT3 knockout mice exposed to vehicle (VEH), a single cocaine injection (SC) or repeated cocaine injections (RC) injections. (**F**) Wild-type mice exposed to RC had significantly greater Zn^2+^ levels compared to VEH-treated WT mice. (**G**) PET experimental timeline and representative horizontal ^65^Zn PET/CT images from wild-type and ZnT3 knockout mice scanned at 14 days after ^65^ZnCl_2_ administration. (**H**) ^65^ZnCl_2_ brain uptake expressed as mean standard uptake value (mSUV) in wild-type and knockout mice scanned at 3, 7 and 14 days after injection. (**I**) Representative ^65^ZnCl_2_ autoradiograms from wild-type and ZnT3 knockout mice scanned using PET at day 15 after ^65^ZnCl_2_ administration and then killed for postmortem verification. (**J**) Statistical contrasts from voxel-wise statistical parametric mapping analyses from vehicle- or cocaine-treated mice exposed to ^65^ZnCl_2_ PET imaging. ppm – parts per million. *p≤0.05, **p≤0.01, ***p≤0.001. All data expressed as Mean ±SEM.

ZnT3 KO mice lack the ability to package Zn^2+^ into synaptic vesicles and by extension cannot release synaptic Zn^2+^ (10). We hypothesized that the increase in cocaine-induced striatal Zn^2+^ we observed in mice was mediated via ZnT3. To assess this, we first confirmed that ZnT3 KO mice lack histochemically-reactive (52) synaptic Zn^2+^. In agreement with prior findings (53), we found that synaptic Zn^2+^ was enriched in discrete areas including medial cortical regions, dorsomedial CPu, and medial NAc in wild-type (WT) mice (Figure 2C) and as expected, ZnT3 KO mice completely lacked synaptic Zn^2+^ (Figure 2D). We then exposed WT and ZnT3 KO mice to daily vehicle (n=6 WT, n=9 KO), a single (20 mg/kg/day, IP) cocaine injection (SC) (n=4 WT, n=4 KO) or repeated cocaine (RC) (20 mg/kg/day for 8 days, IP) (n=5 WT, n=4 KO) injections and killed them 24 hours later followed by dissection of medial frontal cortex and striatum and assessment of Zn^2+^ using TXRF. Vehicle-treated ZnT3 KO mice had significantly lower Zn^2+^ than vehicle-treated WT mice in cortex (Figure S3), where ZnT3 expression and synaptic Zn^2+^ pools are high (Figure 2C). In contrast, vehicle-treated WT and ZnT3 KO mice did not differ in striatal Zn^2+^, as synaptic Zn^2+^ levels in this region are low (Figure 2, E and F) and differences under these basal circumstances were below the detection limit of TXRF (Figure 2E). WT mice exposed to a SC injection did not differ in striatal Zn^2+^ compared to vehicle-treated WT mice [shared control (WT VEH) comparison; unpaired mean difference: vehicle vs. SC=1.52; (95.0% CI −0.31, 2.84), p=0.07] (Figure 2, E and F). In agreement with results from our μSXRF experiments (Figure 2, A and B), WT mice exposed to RC injections had significantly greater striatal Zn^2+^ content than vehicle-treated WT mice [unpaired mean difference: vehicle vs. RC=3.41; (95.0% CI 0.76, 6.87), p=0.03] (Figure 2, E and F). In contrast, cocaine exposure did not lead to any differences in striatal Zn^2+^ in ZnT3 KO mice (Figure 2, E and F).

As a trace element, Zn^2+^ is challenging to study *in vivo*. Consequently, *in vivo* brain Zn^2+^ kinetics have not been previously described. However, *ex vivo* studies using radioactive ^65^ZnCl_2_ have shown that Zn^2+^ has unique brain kinetics, uptake, and biodistribution profiles. Specifically, systemic injection of ^65^ZnCl_2_ takes about 6 days to reach maximal brain uptake (54). Similarly, another study estimated Zn^2+^ turnover in the brain to be a slow process, taking ~7 days (55). Importantly, the *ex vivo* brain distribution of systemically-administered ^65^ZnCl_2_ is dependent on neuronal activity (56) and overlaps with both distribution of ZnT3 protein and histochemically-reactive Zn^2+^ (54), suggesting that the majority of Zn^2+^ uptake to the brain is in the form of synaptic Zn^2+^. In agreement with this, studies have shown that ^65^ZnCl_2_ is transported throughout the brain in a trans-synaptic manner (57). Finally, like synaptic Zn^2+^ (31), ^65^ZnCl_2_ brain uptake is dependent on dietary Zn^2+^ intake (58). Collectively, these studies indicate that ^65^ZnCl_2_ brain uptake can be used as a measure of synaptic Zn^2+^ turnover/metabolism. Interestingly, ^65^Zn produces annihilation photons at 511 keV at a low (~3%) abundance and has a physical half-life of ~244 days. We reasoned that these physical characteristics would be sufficient for noninvasive and longitudinal measurements of ^65^ZnCl_2_ uptake and kinetics using positron emission tomography (PET) and thus could provide *in vivo* assessments of brain Zn^2+^ turnover/metabolism. To confirm, WT (n=2) and ZnT3 KO (n=2) mice were injected intravenously (IV) with 2 μCi/g ^65^ZnCl_2_ and scanned using PET. Owing to the slow kinetics and long biological and physical half-lives of Zn^2+^ (54), mice were scanned longitudinally at different days after ^65^ZnCl_2_ injection (Figure 2G). ^65^Zn showed rapid brain uptake, exhibited slow brain clearance, and was detected in the brain up to 14 days after injection (Figure 2, G and H and Figure S4). ZnT3 KO mice had lower brain uptake of ^65^ZnCl_2_ at 7- and 14-days post injection compared to WT mice (Figure 2H). This was confirmed *ex vivo* using autoradiography, which showed that ^65^Zn brain distribution overlapped with distribution of synaptic Zn^2+^ and ZnT3 protein in WT mice (Figure 2I and Figure S5). Notably, whereas ZnT3 KO mice did show some minimal ^65^ZnCl_2_ uptake, perhaps representing non-specific uptake or a result of other zinc transporters expressed in the brain (59), the distribution of ^65^ZnCl_2_ uptake in KO mice was largely absent from ZnT3-rich regions, indicating that it did not represent the synaptic Zn^2+^ pool. Next, to examine effects of cocaine on ^65^ZnCl_2_ brain uptake, we injected mice with ^65^ZnCl_2_ as above followed by daily vehicle (n=4) or cocaine (n=3) injections (20 mg/kg/day, IP, 8 days) and performed PET. Compared to vehicle, cocaine significantly (1-way anova, treatment main effect, t contrast (1, 4)=2.13; p=0.049) increased uptake of ^65^ZnCl_2_ in brain regions with high ZnT3 and synaptic Zn^2+^ levels, including the prelimbic, cingulate, sensory cortices, NAc, hippocampus, and amygdala but decreased it in areas of low ZnT3 and synaptic Zn^2+^ levels like thalamus (Figure 2J).

### Synaptic Zn^2+^ release increases the *in vivo* potency of cocaine on striatal DA neurotransmission

*In vitro* studies indicate that Zn^2+^ binds to the DAT and causes i) DA reuptake inhibition and ii) increased cocaine DAT binding (18, 21, 22, 60, 61). We reproduced these findings using mouse striatal membranes (n=6 mice/pooled) whereby a physiologically achievable concentration (10 μM) of Zn^2+^ (1) significantly increased [unpaired mean difference: 0 vs. 10 μM =-0.627; (95.0% CI −0.86, −0.41), p<0.001] cocaine binding and affinity to the DAT (Figure 3, A and B, Figure S6). A higher, supraphysiological concentration (1 mM), decreased [unpaired mean difference: 0 vs. 1000 μM =2.51; (95.0% CI 0.88, 3.43), p=0.06] cocaine affinity (Figure 3, A and B). In agreement with the above results, 10 μM Zn^2+^ also significantly increased binding of [^3^H]WIN35,428 to the DAT in tissue sections (n=4 mice) (unpaired t-test, t=4.03; p=0.007) (Figure 3C).

**Figure 3.**
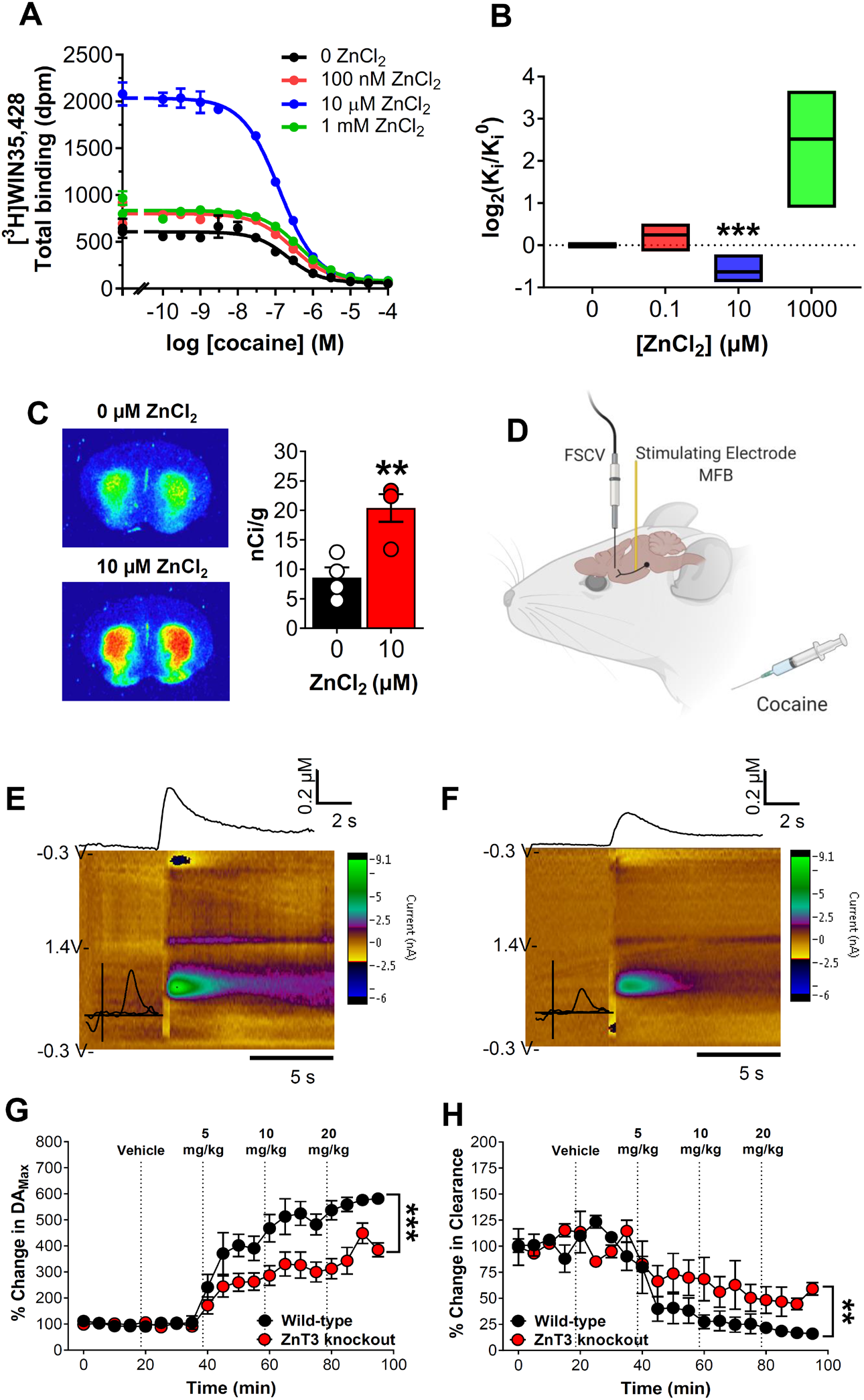
Synaptic Zn^2+^ release increases the *in vivo* potency of cocaine on striatal DA neurotransmission. (**A**) Competition binding curves of cocaine and ZnCl_2_ against [^3^H]WIN-35,428 in mouse striatal tissue. (**B**) 10 μM ZnCl_2_ significantly increased and 1 mM ZnCl_2_ decreased affinity of cocaine in mouse striatum (Ki (±SD)) values in nM; Cocaine: 0 μM Zn, 63±39; 0.1 μM Zn, 77±57; 10 μM Zn, 43±31; 1000 μM Zn, 611±843 and WIN35,428: 0 μM Zn, 9±1.2; 0.1 μM Zn, 9.2±0.8; 10 μM Zn, 5.1±1.1; 1000 μM Zn, 13.4±1.8). (**C**) Autoradiograms at the level of mouse striatum showing that 10 μM ZnCl_2_ significantly increased [^3^H]WIN-35,428 binding. (**D**) Experimental design of fast scan cyclic voltammetry (FSCV) experiment. (**E**) Representative FSCV color plots from wild-type and (**F**) ZnT3 knockout mice showing changes in electrically-evoked dopamine (DA) after a 10 mg/kg IP cocaine injection. (**G**) FSCV time-course plots showing significantly lower percent change in electrically-evoked DA_Max_ and (**H**) faster DA Clearance rate in ZnT3 knockout compared to wild-type mice as a function of vehicle or escalating IP cocaine injections. *p≤0.05, **p≤0.01, ***p≤0.001. All data expressed as Mean ±SEM.

To assess cocaine’s *in vivo* effects at the DAT as a function of synaptic Zn^2+^, we performed *in vivo* fast scan cyclic voltammetry (FSCV) in the striatum of WT (n=4) and ZnT3 KO (n=4) mice. Electrical stimulation of the medial forebrain bundle (MFB) was used to evoke DA release in the striatum followed by escalating cocaine injections (5, 10, 20 mg/kg, IP) (Figure 3D). WT and ZnT3 KO mice did not differ in baseline DA release or clearance (Figure S7). <u>DA release:</u> Compared to Vehicle, we found that 5, 10, and 20 mg/kg cocaine significantly increased electrically-evoked extracellular DA release in WT mice, but ZnT3 KO mice only showed a significant increase at 20 mg/kg cocaine (Figure S7). ZnT3 KO mice also showed significantly lower electrically-evoked extracellular DA release compared to WT mice (2-way RM anova; genotype x time interaction, F(19, 114)=5.46; p<0.001) (Figure 3E, G). <u>DA clearance:</u> Compared to Vehicle, 5, 10, and 20 mg/kg cocaine significantly decreased DA clearance in WT mice as compared to Vehicle, but ZnT3 KO mice only showed significantly lower DA clearance at 20 mg/kg cocaine (Figure S7). ZnT3 KO mice showed significantly greater DA clearance in response to cocaine compared to WT mice (2-way RM anova; genotype x time interaction, F(19, 114)=2.35 p=0.0029) (Figure 3F, H).

### Synaptic Zn^2+^ release potentiates cocaine locomotor sensitization, reward, seeking and is required for cocaine-induced increases in DAT

We hypothesized that cocaine-induced synaptic Zn^2+^ release would modulate cocaine-related behavior. To examine this, we first tested ZnT3 KO mice for development and expression of cocaine-induced locomotor sensitization. WT (n=15) and ZnT3 KO mice (n=16) both developed locomotor sensitization to daily injections of cocaine (10 mg/kg/day, IP, 5 days) (2-way RM anova with Holm-Sidak multiple comparisons, genotype x session interaction, F(3, 59)=7.07; p=0.004) (Figure 4A). However, ZnT3 KO mice (n=16) showed significantly lower locomotor activity compared to cocaine-treated WT mice on Day 1 (t=3.09; p=0.01) and Day 5 (t=3.22; p=0.009) of the procedure. As expected, WT mice injected with cocaine showed significantly greater locomotor activity than vehicle-treated WT (n=16) mice on Day 1 (t=5.54; p<0.001) and Day 5 (t=8.71; p<0.001). KO mice injected with cocaine showed significantly greater locomotor activity than vehicle-treated KO (n=16) mice on Day 5 (t=5.68; p<0.001) but not on Day 1 (t=2.48; p=0.08). One week later, vehicle-treated and cocaine-treated mice were tested for expression of cocaine locomotor sensitization via exposure to escalating cocaine injections (5, 10, and 20 mg/kg, IP). WT mice with prior cocaine exposure (cocaine-treated) (n=9) showed significantly greater (2-way RM anova with Holm-Sidak multiple comparisons, genotype x time interaction, F(79, 1343)=1.85; p<0.001) expression of cocaine locomotor sensitization at 5 (66 min; t=3.65; p=0.02, 69 min; t=3.72; p=0.01) and 10 mg/kg cocaine (123 min; t=5.24; p<0.001, 126 min; t=6.03; p<0.001, 129 min; t=5.19; p<0.001, 132 min; t=3.75; p=0.01, 135 min; t=3.54; p=0.03) compared to vehicle-treated wild-type mice (n=10) (Figure 4B). In contrast, cocaine-treated KO mice (n=10) did not show a significant increase in locomotor activity (2-way RM anova, genotype x time interaction, F(79, 1422)=0.9197; p=0.67) compared to vehicle-treated KO mice (n=10) (Figure 4C). Neither WT nor ZnT3 KO cocaine-treated mice differed from the corresponding vehicle-treated mice at a 20 mg/kg dose of cocaine (Figure 4, B and C).

**Figure 4.**
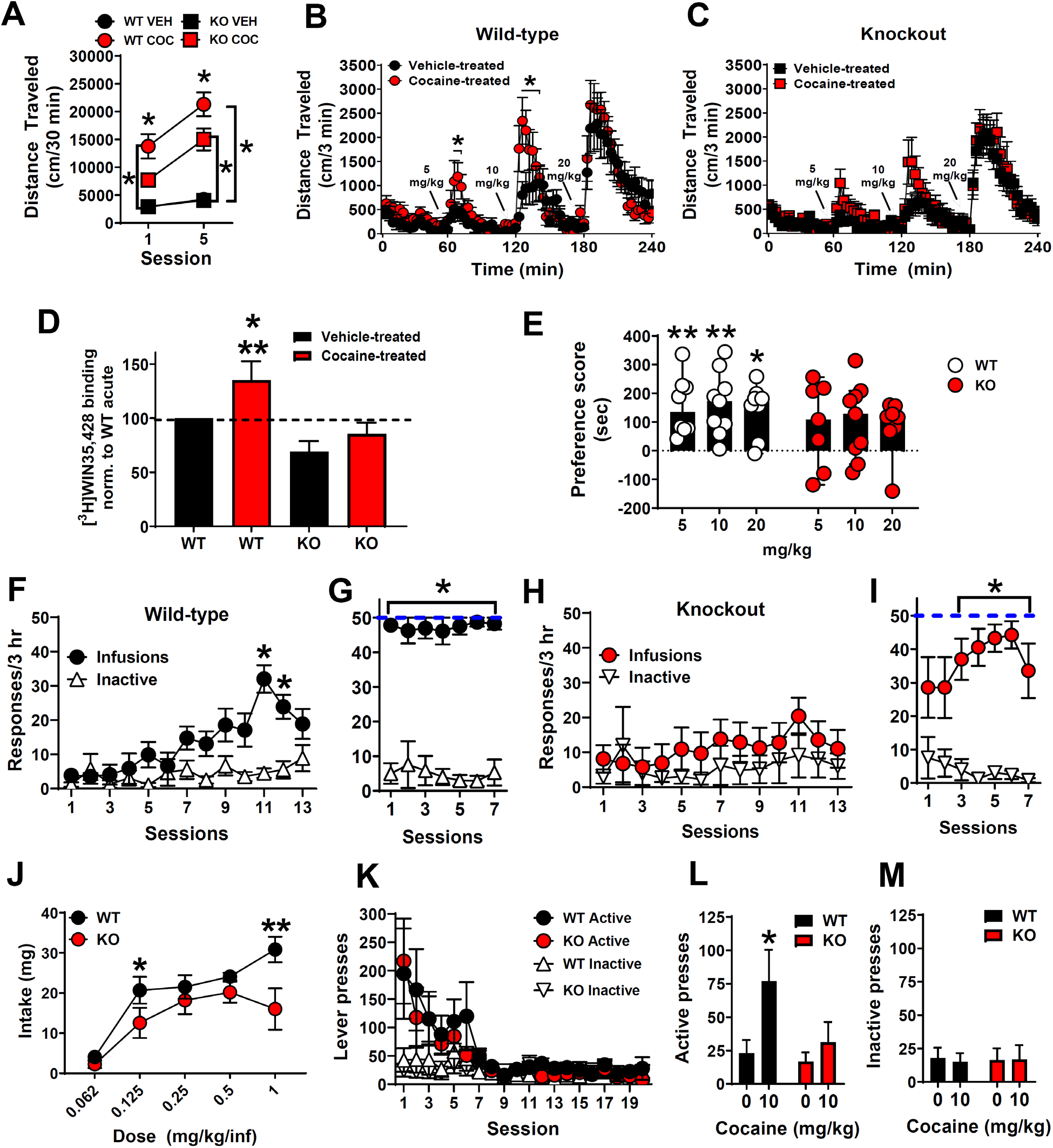
Synaptic Zn^2+^ release potentiates cocaine locomotor sensitization, reward, seeking and is required for cocaine-induced increases in DAT. (**A**) ZnT3 knockout (KO) mice injected with 10 mg/kg cocaine (COC) showed significantly lower locomotor activity compared to cocaine-injected wild-type (WT) mice. Compared to vehicle, cocaine significantly increased locomotor activity in both WT and KO mice on Day 5 but only in WT mice on Day 1. (**B**) Cocaine-treated WT mice showed significantly greater expression of cocaine locomotor sensitization at 5 and 10 mg/kg cocaine compared to vehicle-treated WT mice. (**C**) Cocaine-treated ZnT3 KO mice do not differ in expression of cocaine locomotor sensitization compared to vehicle-treated KO mice. (**D**) Cocaine-treated WT mice had significantly greater DAT binding than cocaine-treated ZnT3 KO mice and vehicle-treated KO mice and showed a trend toward significantly greater DAT binding than vehicle-treated WT mice. (**E**) WT mice showed significant preference for a chamber paired with cocaine at 5, 10, and 20 mg/kg. ZnT3 KO mice did now show preference for a chamber paired with cocaine at any dose. (**F**) Wild-type mice exposed to a training dose of cocaine (1 mg/kg/inf.) showed significantly greater cocaine-reinforced presses compared to inactive lever presses. (**G**) Once trained, WT mice were exposed to a lower cocaine dose (0.5 mg/kg/inf.), immediately reached the maximum number of presses allowed per session, and showed significantly greater cocaine-reinforced presses compared to inactive lever presses on all sessions. (**H**) ZnT3 knockout mice exposed to a training dose of cocaine (1 mg/kg/inf.) did not show any significant differences between cocaine-reinforced and inactive lever pressing. (**I**) ZnT3 knockout mice exposed to a lower cocaine dose (0.5 mg/kg/inf.) showed significantly greater cocaine-reinforced presses compared to inactive lever presses during the last 5 sessions. (**J**) ZnT3 KO mice showed significantly lower cocaine intake at 1 mg/kg/inf. and at 0.125 mg/kg/inf. compared to WT mice. (**K**) WT and ZnT3 KO mice did not differ in extinction of cocaine self-administration. (**L**) After extinction of cocaine self-administration behavior, WT mice showed reinstatement of cocaine self-administration and significantly greater active lever presses after cocaine priming compared to ZnT3 KO mice. (**N**) WT and ZnT3 KO mice did not differ in inactive lever responding during cocaine-primed reinstatement. *p≤0.05. All data except panel E expressed as Mean ±SEM. Panel E is expressed as Median ±95% CI.

Cocaine exposure increases DAT levels (47, 48, 62) which is thought to serve as a compensatory adaptation to DAT blockade after repeated cocaine exposure. We hypothesized that ZnT3 KO mice, which lack cocaine-induced Zn^2+^ release and expression of cocaine-induced locomotor sensitization, would be insensitive to cocaine-induced increases in striatal DAT. To test this, mice from the above sensitization experiments were killed 24 hours after the last cocaine injection, the striatum was dissected, and DAT binding assays were performed using [^3^H]WIN-35,428 (3 repetitions per curve, in triplicate) (one-way ANOVA: F(3, 8)=6.23; p=0.01). Pairwise comparisons (Holm-Sidak) showed that vehicle-treated WT mice (n=6) did not differ from vehicle- or cocaine-treated ZnT3 KO mice (n=6) in [^3^H]WIN-35,428 binding (Figure 4D). However, cocaine-treated WT mice (n=6) showed a trend toward greater [^3^H]WIN-35,428 binding (t=2.21; p=0.058) compared to vehicle-treated WT mice (n=6) and significantly greater [^3^H]WIN-35,428 binding than both vehicle-treated (n=6) (t=4.14; p=0.009) and cocaine-treated KO mice (n=6) (t=3.12; p=0.02).

Next, we tested whether synaptic Zn^2+^ release is involved in cocaine reward by assessing the extent to which ZnT3 KO mice develop cocaine conditioned place preference (CPP). As expected, we found that WT mice showed significant preference for a chamber paired with cocaine (one sample Wilcoxon (non-parametric) test with theoretical median set to 0) at 5 (n=8) (p=0.007; 95% CI: 41.3 to 336.1), 10 (n=9) (p=0.003; 95% CI: 59.2 to 297.3), and 20 (n=8) (p=0.01; 95% CI: −9.8 to 258.4) mg/kg. In contrast, ZnT3 KO mice did not show significant preference for the chamber paired with cocaine at any dose: 5 mg/kg (n=7) (p=0.21; 95% CI: −118.8 to 256.2), 10 mg/kg (n=9) (p=0.07; 95% CI: −47.2 to 208.5) and 20 mg/kg (n=7) (p=0.10; 95% CI: −140.6 to 159.3) (Figure 4E).

Finally, we examined whether synaptic Zn^2+^ is involved in intravenous cocaine self-administration (SA). WT mice (n=9) showed robust acquisition of cocaine SA (1 mg/kg/infusion for 13 days), as evidenced by significantly greater (Mixed effects anova with Holm-Sidak multiple comparisons); genotype x session interaction, F(12, 190)=6.12; p<0.001) and sustained responding on the cocaine-reinforced active lever over the non-reinforced inactive lever on days 11 and 12 (session 11: t=6.48; p<0.001, session 12: t=4.25; p=0.009) (Figure 4F). In contrast, ZnT3 KO mice (n=9) failed to acquire cocaine SA at the 1 mg/kg/infusion dose (Figure 4H) as they did not show any significant differences (2-way RM anova with Holm-Sidak multiple comparisons, genotype x session interaction, F(12, 192)=1.27; p=0.23) in cocaine-reinforced vs. inactive lever pressing. All mice were then switched to a lower cocaine dose (0.5 mg/kg/infusion for 7 days). WT mice (n=7) immediately reached the maximum allowed number (50) of infusions (Figure 4G) and showed significantly greater cocaine-reinforced presses (2-way RM anova with Holm-Sidak multiple comparisons, lever main effect, F(1, 12)=116.7; p<0.001) compared to inactive lever presses (session 1: t=12.54; p<0.001, session 2: t=5.03; p=0.004, session 3: t=7.91; p<0.001, session 4: t=9.39; p<0.001, session 5: t=15.45; p<0.001, session 6: t=23.33; p<0.001, session 7: t=10.42; p<0.001). At the 0.5 mg/kg/infusion dose, ZnT3 KO mice took longer (3 sessions) to learn to discriminate the cocaine-reinforced active lever over the inactive lever and also took longer (5 sessions) to reach 50 infusions (Figure 4I). Eventually, ZnT3 KO mice (n=7) showed significantly greater cocaine-reinforced presses (2-way RM anova with Holm-Sidak multiple comparisons, lever x time interaction, F(6, 71)=3.9; p=0.002) compared to inactive lever presses (session 3: t=4.74; p=0.007, session 4: t=7.03; p=0.002, session 5: t=8.86; p<0.001, session 6: t=9.82; p<0.001, session 7: t=4.01; p=0.04). After acquisition of cocaine self-administration, mice were tested at lower doses of cocaine. WT and KO mice did not differ in the number of cocaine infusions at these lower doses (Figure S8). However, compared to WT mice (n=6), KO mice (n=5) showed significantly lower cocaine intake (mixed effects RM analysis with Holm-Sidak multiple comparisons, genotype x dose interaction, F(4, 44) =2.92) at 1 mg/kg/inf. (t=3.26; p=0.001) and at 0.125 mg/kg/inf. (t=2.03; p=0.04) doses (Figure 4J). Mice were then assessed for extinction of cocaine self-administration, but no genotype differences were observed (Figure 4K). After mice had extinguished their lever responding for cocaine, they were tested for reinstatement (relapse) responding to a cocaine priming injection (10 mg/kg, IP). WT mice (n=5) showed reinstatement of cocaine self-administration and significantly greater active lever presses (2-way RM anova with Holm-Sidak multiple comparisons, genotype x dose interaction, F(1, 10)=5.09; p=0.04) after cocaine priming (t=3.17; p=0.005) compared to ZnT3 KO mice (n=5), which did not reinstate (Figure 4L and M). WT mice (n=5) and ZnT3 KO mice (n=5) did not differ in inactive lever responding (2-way RM anova with Holm-Sidak multiple comparisons, genotype x dose interaction, F(1, 10)=0.17; p=0.68) during cocaine-primed reinstatement.

### Dietary Zn^2+^ deficiency decreases brain Zn^2+^ and attenuates cocaine locomotor sensitization, reward, and striatal DAT binding

Synaptic Zn^2+^ levels and ZnT3 expression are modulated by dietary Zn^2+^ intake (30, 31). Chronic drug abuse (including cocaine) in humans is associated with malnutrition and dysregulated peripheral Zn^2+^ levels (33–38). However, whether dietary Zn^2+^ availability can modulate the behavioral effects of cocaine has not been previously reported.

To examine this, we fed mice, for approximately one month, diets that were custom-formulated by a commercial vendor to contain either 30 (considered an adequate level of Zn^2+^ intake) or 5 (a low amount of Zn^2+^ intake) ppm Zn^2+^. We first verified the Zn^2+^ content of each diet using TXRF. According to our measurements, the 30 ppm diet contained ~28 ppm Zn^2+^ and the 5 ppm diet contained ~8 ppm Zn^2+^ (Figure 5A) (we will however refer to them here as 30 and 5 ppm Zn^2+^ diets to remain consistent with prior studies that have used similar formulations). Mice fed a 5 ppm Zn^2+^ diet (n=32) did not differ in body weight from mice fed a 30 ppm Zn^2+^ diet (n=32) (Figure 5B). Mice fed a 5 ppm Zn^2+^ diet (n=32) showed a significant decrease (2-way RM anova with Holm-Sidak multiple comparisons; genotype x time interaction, F(15, 930)=10.07; p<0.001) in food intake 3 days (t=11.37; p<0.001) after diet initiation compared to mice fed a 30 ppm diet (n=32) though soon afterwards normalized to the same levels of intake (Figure 5C). The low Zn^2+^ diet was effective in decreasing brain Zn^2+^ content as mice fed a 5 ppm Zn^2+^ diet (n=7) showed significantly lower Zn^2+^ levels in frontal cortex (unpaired t-test; t=3.15; p=0.01) compared to mice fed a 30 pm Zn^2+^ diet (n=5) as assessed via TXRF (Figure 5D).

**Figure 5.**
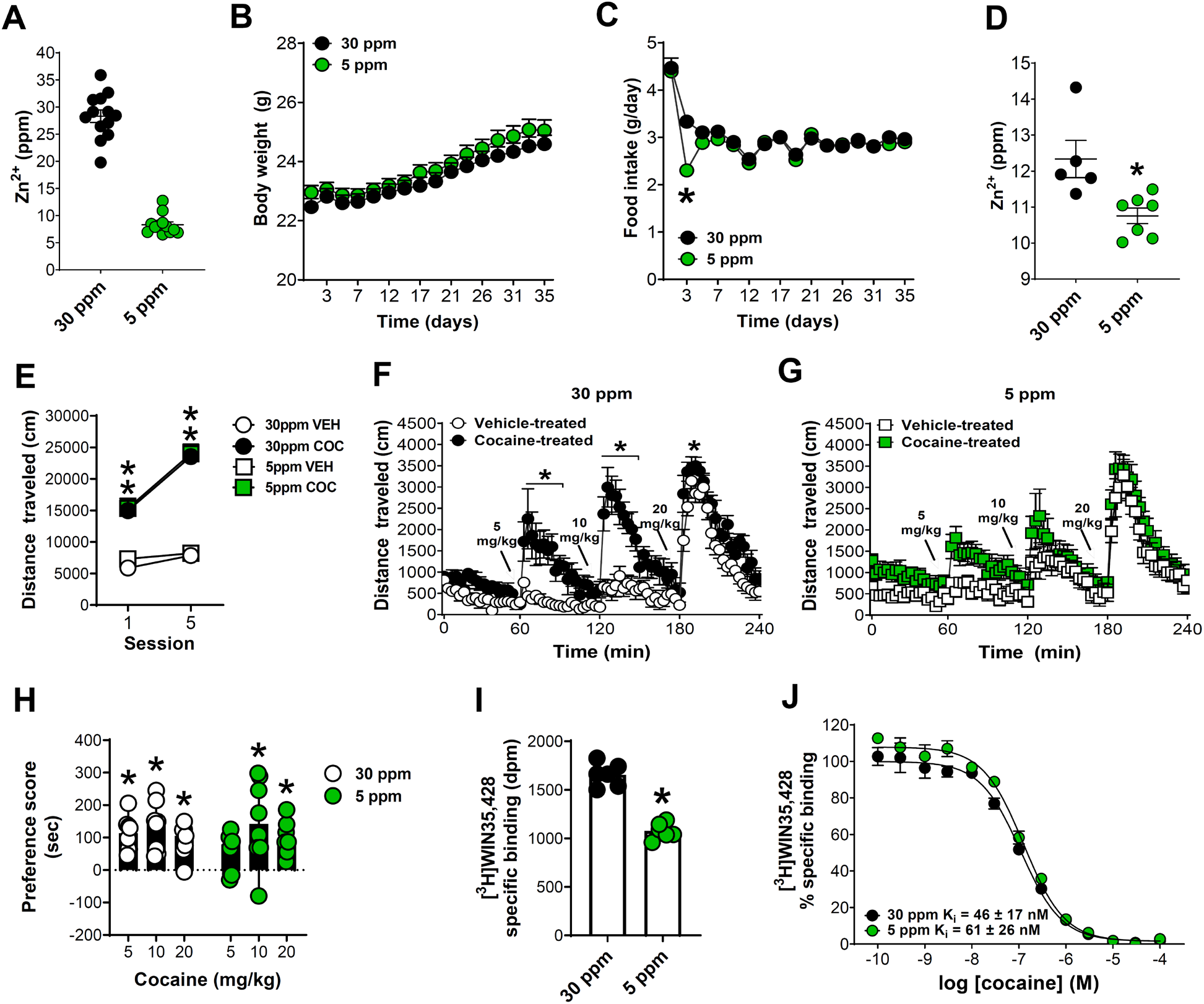
Dietary Zn^2+^ deficiency decreases brain Zn^2+^ and attenuates cocaine locomotor sensitization, reward and striatal DAT binding. (**A**) Total reflection X-ray spectroscopy (TXRF)-based verification of Zn^2+^ content in custom-made diets. (**B**) Mice fed a 5 ppm Zn^2+^ diet did not differ in body weight from mice fed a 30 ppm Zn^2+^ diet. (**C**) Mice fed a 5 ppm Zn^2+^ diet showed a significant decrease in food intake 3 days after diet initiation compared to mice fed a 30 ppm diet but did not differ at other times. (**D**) Mice fed a 5 ppm Zn^2+^ diet showed significantly lower total Zn^2+^ in frontal/cingulate cortex compared to mice fed a 30 pm Zn^2+^ diet. (**E**) Mice fed 5 ppm or 30 ppm Zn^2+^ diets did not differ in development of cocaine (COC) locomotor sensitization (VEH-vehicle). (**F**) Cocaine-treated 30 ppm diet mice (n=6) showed significantly greater expression of cocaine locomotor sensitization at 5, 10 mg/kg and 20 mg/kg cocaine compared to vehicle-treated 30 ppm mice (n=13). (**G**) Cocaine-treated 5 ppm mice did not show any significant difference in expression of locomotor sensitization compared to vehicle-treated 5 ppm mice. (**H**) Mice fed a 30 ppm Zn^2+^ diet showed significant preference for a chamber paired with cocaine at 5, 10, and 20 mg/kg. Mice fed a 5 ppm Zn^2+^ diet showed significant preference for a chamber paired with cocaine at 10, and 20 mg/kg but not at 5 mg/kg. (**I**) Mice exposed to cocaine and a 5 ppm Zn^2+^ diet showed significantly lower DAT binding in striatum and (**J**) a mild decrease in cocaine affinity compared to mice exposed to cocaine and a 30 ppm diet. dpm - disintegrations per minute. *p≤0.05. All data except panel H expressed as Mean ±SEM. Panel H is expressed as Median ±95% CI.

Next, we performed cocaine locomotor sensitization in 30 ppm and 5 ppm mice using the same procedures as in the above ZnT3 KO experiments. Mice fed 5 ppm or 30 ppm Zn^2+^ diets did not differ in the development of cocaine locomotor sensitization (Figure 5E). 30 ppm diet-fed mice injected with cocaine (n=14) showed significantly greater locomotor activity (2-way RM anova with Holm-Sidak multiple comparisons; genotype x session interaction, F(3, 52)=23.57; p<0.001) than 30 ppm diet-fed mice injected with vehicle (n=14) at Day 1 (t=8.29; p<0.001) and Day 5 (t=14.27; p<0.001). Similarly, 5 ppm mice injected with cocaine (n=14) showed significantly greater locomotor activity compared to 5 ppm mice injected with vehicle (n=14) at Day 1 (t=6.95; p<0.001) and Day 5 (t=13.95; p<0.001). 30 ppm mice injected with cocaine also showed significantly greater locomotor activity at Day 5 than at Day 1 (t=9.98; p<0.001) and 5 ppm mice injected with cocaine also showed significantly greater locomotor activity at Day 5 than at Day 1 (t=10.06; p<0.001) (Figure 5E). Seven days later, mice were examined for expression of cocaine locomotor sensitization. We found that cocaine-treated mice fed the 30 ppm diet (n=6) showed significantly greater expression of cocaine locomotor sensitization (2-way RM anova with Holm-Sidak multiple comparisons; genotype x time interaction, F(79, 790)=3.98; p<0.001) at 5 (66 min; t=5.69; p<0.001, 69 min; t=4.68; p<0.001, 72 min; t=3.76; p=0.01, 75 min; t=3.78; p=0.01, 78 min; t=4.29; p=0.001, 81 min; t=3.79; p=0.01, 84 min; t=4.23; p=0.002), 10 mg/kg (123 min; t=5.37; p<0.001, 126 min; t=7.42; p<0.001, 129 min; t=6.99; p<0.001, 132 min; t=6.58; p=0.01, 135 min; t=5.08; p=0.03, 138 min; t=5.36; p=0.03, 141 min; t=4.39; p=0.03, 144 min; t=4.14; p=0.03, 147 min; t=3.78; p=0.03) and 20 mg/kg cocaine (183 min; t=3.43; p=0.04) compared to vehicle-treated mice fed the same 30 ppm diet (n=13) (Figure 5F). In contrast, cocaine-treated mice fed the 5 ppm diet (n=6) did not show a significant difference in expression of locomotor sensitization compared to vehicle-treated mice fed the 5 ppm diet (n=6) (2-way RM anova with Holm-Sidak multiple comparisons; genotype x time interaction, F(79, 790)=0.9342; p=0.64) (Figure 5G).

Next, we tested whether mice fed 30 ppm and 5 ppm Zn^2+^ diets differed in cocaine preference using the same CPP procedure as in ZnT3 KO mice. Mice fed the 30 ppm Zn^2+^ diet showed significant preference (one sample Wilcoxon test with theoretical median set to 0) for the chamber paired with cocaine at 5 mg/kg (n=8) (p=0.007; 95% CI: 45.8 to 205.5), 10 mg/kg (n=8) (p=0.007; 95% CI: 42.6 to 245.4), and 20 mg/kg (n=7) (p=0.003; 95% CI: −6.0 to 149.6). Mice fed a 5 ppm Zn^2+^ diet showed significant preference (one sample Wilcoxon test with theoretical median set to 0) for the chamber paired with cocaine at 10 mg/kg (n=8) (p=0.039; 95% CI: −79.8 to 298.0), and 20 mg/kg (n=8) (p=0.007; 95% CI: 26.2 to 185.9) but not at 5 mg/kg (n=8) (p=0.10; 95% CI: −30.1 to 125.7) (Figure 5H).

Finally, we examined whether dietary Zn^2+^ availability had any effects on DAT binding in the striatum. Mice fed 30 ppm and 5 ppm Zn^2+^ diets and exposed to escalating cocaine injections (5, 10, and 20 mg/kg, IP, one injection/hour) were killed 24 hours after the last injection and their brains were assessed for striatal [^3^H]WIN-35,428 binding. We found that mice exposed to cocaine and a 5 ppm Zn^2+^ diet (n=6) showed significantly lower striatal DAT binding (Figure 5I) (unpaired t-test; t=9.63; p<0.001) (3 repetitions per group, in triplicate) and a decrease in cocaine affinity compared to mice exposed to cocaine and a 30 ppm Zn^2+^ diet (n=6) (5ppm Ki=61 ± 26 nM; 30 ppm Ki=46 ± 17 nM) (3 repetitions per curve, in triplicate) (Figure 5J).

## Discussion

Here we show that human subjects who died of cocaine overdose had significantly lower Zn^2+^ content in the striatum (caudate) compared to matched controls. We also show that striatal Zn^2+^ levels in these subjects negatively correlated with plasma levels of benzoylecgonine, indicating that low striatal Zn^2+^ levels were associated with greater cocaine intake.

To further explore the relationship between cocaine exposure and striatal Zn^2+^, we performed studies using mice lacking ZnT3, a neuronal Zn^2+^ transporter necessary for synaptic Zn^2+^ release. We found that repeated cocaine injections increased striatal Zn^2+^ content (in CPu and NAc) in normal mice but not in mice lacking ZnT3, indicating that cocaine exposure increases synaptic Zn^2+^ levels in the striatum via the actions of ZnT3. We also found that repeated cocaine injections increased Zn^2+^ uptake in regions with high synaptic Zn^2+^ and ZnT3 content, including the NAc indicating that cocaine exposure increases synaptic Zn^2+^ turnover/metabolism in these regions.

We hypothesized that cocaine-induced increases in synaptic Zn^2+^ release would be relevant to cocaine’s *in vivo* actions at the DAT, given the known *in vitro* interactions between Zn^2+^, DAT, and cocaine. Using *in vitro* assays, we first confirmed that a physiologically-relevant concentration of Zn^2+^ increased the ability of cocaine to bind to DAT. Then, to extend the relevance of this finding to an *in vivo* context, we used *in vivo* FSCV and found that deletion of ZnT3, and hence, loss of synaptic Zn^2+^ release, attenuated cocaine’s effects on striatal DA neurotransmission. Consistent with this observation, mice lacking ZnT3 were insensitive to cocaine-induced increases in striatal DAT binding and showed attenuated behavioral responses to cocaine in several procedures such as locomotor sensitization, CPP, and IV self-administration, indicating that synaptic Zn^2+^ promotes cocaine sensitization, reward, and cocaine-seeking behavior.

Zn^2+^ is obtained almost exclusively from the diet. Taking our above findings into account, we reasoned that low environmental availability of Zn^2+^ and specifically, reduced dietary Zn^2+^ intake, would attenuate behavioral responses to cocaine. As predicted, we found that mice exposed to a diet with low Zn^2+^ content showed lower brain Zn^2+^ levels, decreased cocaine-induced increases in striatal DAT binding, and attenuated behavioral responses to cocaine compared to mice fed a diet with adequate Zn^2+^ content. This indicates that low dietary Zn^2+^ intake decreases brain Zn^2+^ levels, and attenuates cocaine locomotor sensitization, cocaine preference, and cocaine-induced increases in striatal DAT.

Taken together, our above findings suggest that synaptic Zn^2+^ release in the striatum plays a critical role in cocaine’s effects on striatal DA neurotransmission and consequently in the neurobiological and behavioral adaptations associated with cocaine exposure. A summary depicting this proposed mechanism is shown in Figure 6.

**Figure 6.**
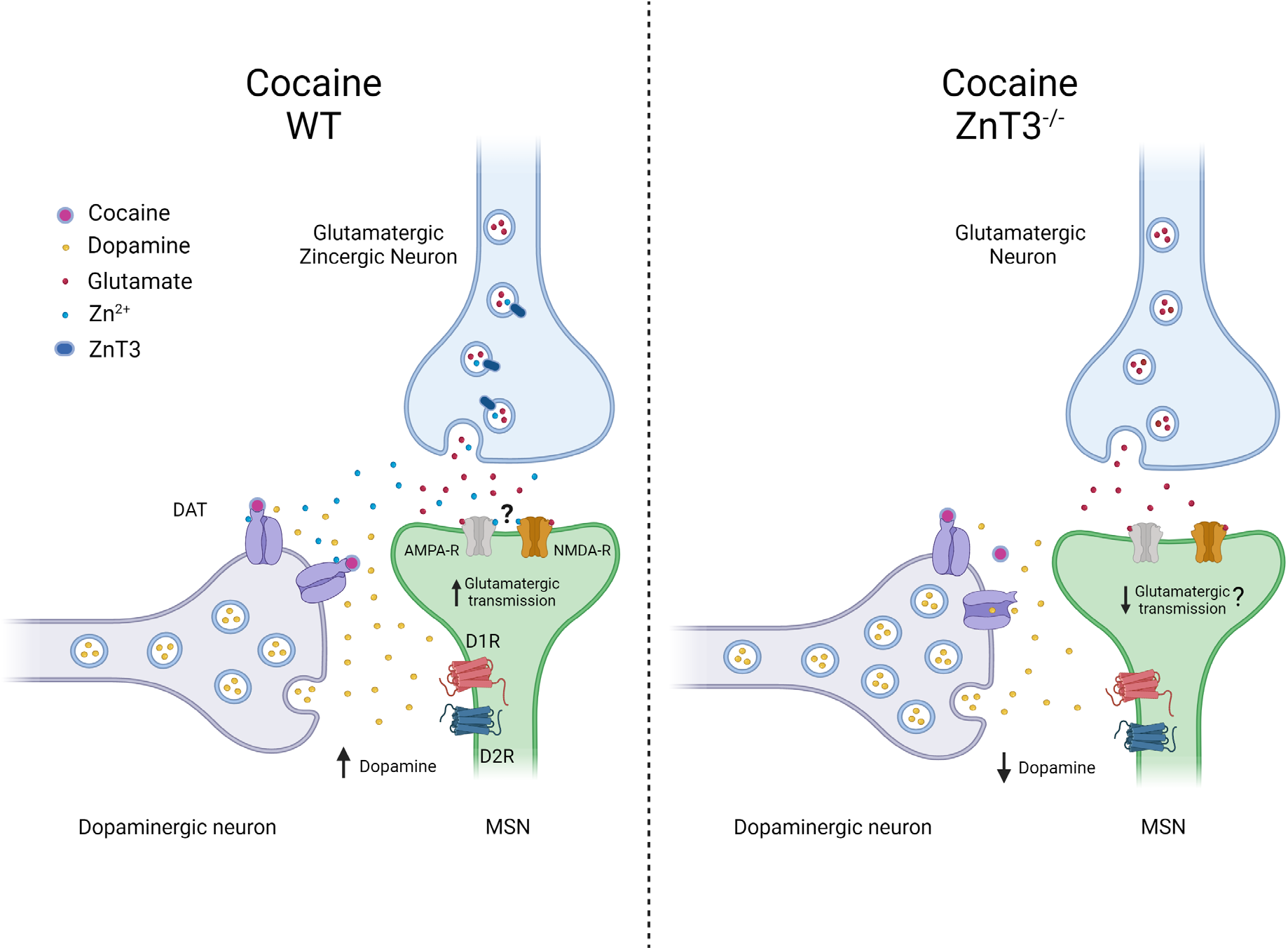
Schematic representation of the putative mechanism by which presynaptic Zn^2+^ modulates the effects of cocaine on dopamine and glutamate neurotransmission in the striatum in wild-type (WT) and ZnT3 knockout mice. The image on the left shows the effect of synaptic Zn^2+^ on increasing cocaine’s efficacy by binding to the DAT on dopaminergic neurons. It also acknowledges the putative effects of Zn^2+^ on glutamate neurotransmission in the context of cocaine via its interaction with AMPA and NMDA receptors. The image on the right illustrates the effect of ZnT3 deletion, and consequently the loss of presynaptic Zn^2+^ release, in attenuating the effects of cocaine on striatal dopamine and glutamate neurotransmission.

The results from our experiments in mice suggest that the Zn^2+^ deficits we identified in humans who died from cocaine overdose may arise from a combination of inadequate nutrition (i.e. low dietary Zn^2+^ intake) and increased Zn^2+^ turnover/metabolism brought upon by chronic cocaine use, although we cannot rule out individual or combined contributions of diet, cocaine use, or use of other drugs/substances in these effects. The negative correlation between striatal Zn^2+^ content and plasma benzoylecgonine levels in postmortem human samples, taken together with the rest of our data, may reflect the notion that individuals with inherently low striatal Zn^2+^ levels would be less sensitive to the effects of cocaine and therefore would need to compensate by consuming more of the drug. This would implicate Zn^2+^ as an environmental factor that could influence vulnerability to the effects of cocaine and potentially to the development of cocaine use disorders or addiction.

Our results have important implications for both general DA-dependent behaviors and especially for the prevention and treatment of cocaine addiction. Specifically, our findings suggest that dietary Zn^2+^ intake, and potentially, impaired Zn^2+^ absorption or excretion mechanisms, are implicated in cocaine reward, seeking, and relapse. Consequently, we suggest that the Zn^2+^ status of patients with cocaine addiction should be taken into consideration, especially since Zn^2+^ deficiency varies in prevalence across social demographics and is found in higher proportion in developing countries (32).

Our findings also expand the current understanding of cocaine’s neurobiological effects. DA and glutamate systems converge at the level of the striatum to modulate behaviors that influence cocaine seeking and abuse (46, 63). Synaptic Zn^2+^, mediated by ZnT3, is packaged along with glutamate in the same synaptic vesicles and is released by glutamatergic terminals in the striatum upon neuronal activation. As such, the cocaine-induced synaptic Zn^2+^ increases we report here would be expected to accompany the well-described increase in synaptic glutamate neurotransmission in the striatum following repeated cocaine exposure (45, 46, 64, 65). Indeed, cocaine-induced increases in synaptic glutamate are thought to result, in part, from reductions in extrasynaptic glutamate arising from decreased cystine-glutamate exchange in response to repeated cocaine use (63, 66). Specifically, cocaine-induced reductions in extrasynaptic glutamate weaken signaling at presynaptic metabotropic glutamate receptors, thereby increasing synaptic glutamate levels (64) and resulting in excitatory synaptic strengthening. This mechanism may underlie the increases in cocaine-induced synaptic Zn^2+^ that we observe.

Zn^2+^ binds to NMDA and AMPA receptors (67, 68) and it is likely that, in addition to affecting DA neurotransmission via binding to DAT, cocaine-induced increases in synaptic Zn^2+^ release may also exert direct allosteric interactions at ionotropic glutamate receptors to influence cocaine-dependent glutamate neurotransmission, synaptic plasticity, and behaviors such as cocaine locomotor sensitization and cocaine priming-induced reinstatement of cocaine seeking (45, 46, 65, 69). However, the precise mechanism for how cocaine-induced Zn^2+^ release would lead to such effects is complicated by the fact that Zn^2+^ can exert bidirectional effects at AMPA receptors (70), it can inhibit both synaptic and extrasynaptic NMDA receptors (68), and finally, that cocaine induces projection- and cell type-specific adaptations in the NAc (71), that not all NAc glutamatergic synapses release Zn^2+^, and that the specific zincergic synapses, and the postsynaptic cells they target in regions such as the NAc have not yet been defined. For all these reasons, the specific effects of cocaine-induced Zn^2+^ release on glutamatergic signaling are unclear and require further investigation.

Another possibility may be that Zn^2+^ interacts with cystine-glutamate exchange mechanisms directly to produce changes in cocaine behaviors. Indeed, we found here that ZnT3 KO mice failed to reinstate cocaine-primed self-administration of cocaine, suggesting that synaptic Zn^2+^ release is necessary for cocaine-primed seeking. N-acetylcysteine (NAC) is a derivative of the amino acid cysteine and has been proposed as a treatment for substance use disorders (72). NAC prevents cocaine-primed reinstatement of drug seeking after cocaine self-administration by stimulating cystine-glutamate exchange (63). Importantly, NAC also has metal-chelating capabilities (Zn^2+^ binds with high affinity to cysteines (73)) and systemic administration of NAC in mice has been shown to decrease peripheral Zn^2+^ levels (74). Accordingly, it may be possible that in addition to the effects of NAC in decreasing cocaine-primed seeking via activation of cystine-glutamate exchange, its effects may also be driven, in part, via direct interaction with synaptic Zn^2+^. In this case, NAC would effectively act as a Zn^2+^ chelating agent, decreasing synaptic Zn^2+^ levels, which would attenuate cocaine’s effects at the DAT and decrease cocaine-seeking. However, this could also occur concomitantly to NAC’s activation of cystine-glutamate exchange, which would increase extrasynaptic glutamate to restore presynaptic metabotropic glutamate receptor tone and thereby reduce synaptic glutamate and potentially, co-released Zn^2+^.

In addition to cocaine, other psychostimulants interact with the DAT to produce their effects of behavior. We propose that the cocaine-dependent changes in Zn^2+^ that we describe here are not specific to cocaine but may also be elicited by other psychostimulants that modulate DAT function and increase corticostriatal glutamate neurotransmission (45, 46, 75), and potentially other drugs of abuse, including alcohol (76) which is strongly associated with Zn^2+^ deficiency (76).

Finally, in addition to its critical role in addiction, the DAT is the primary molecular target for stimulant medications used in the treatment of attention-deficit hyperactivity disorder (ADHD). Studies indicate that medication response is reduced in Zn^2+^-deficient ADHD patients (77). Our findings here offer a mechanistic explanation for these clinical observations and suggest that Zn^2+^ supplementation may improve the efficacy of stimulant-based ADHD medications.

In conclusion, our findings expand current knowledge regarding the direct pharmacological mechanism of action of cocaine and the neurobiological mechanisms involved in cocaine’s behavioral effects by identifying a critical role for the trace element Zn^2+^ as an environmentally derived modulator of cocaine affinity, potency, plasticity, reward, and seeking.

## Acknowledgments and Author contributions

JLG, JB, SL, DM, KW, MLC, LAR, RJE, CJ, GB, JK, MP, MB, ZX, GT, OS, and MM designed and/or performed experiments and/or analyzed data. ZX, GT, and MM supervised work. JLG, JB, and MM wrote the paper with input from all coauthors. The authors thank Dr. Thanos Tzounopoulos (University of Pittsburgh) for sharing ZnT3 knockout mice, Dr Richard Dyck (University of Calgary) for advice regarding Zn^2+^ staining, Dr. Yavin Shaham (NIDA) for experimental advice in behavioral experimentation and for manuscript comments, and Dr. Marisela Morales (NIDA), Dr. Antonio Lanzirotti, Dr. Keith Jones and William Rao at the National Synchrotron Light Source (NSLS) X26a Beamline at Brookhaven National Laboratory for instrumentation access. This work was supported by the NIDA Intramural Research Program (DA000069), the NIDA Medication Development Program (DA000611) and the Department of Energy (DOE) GeoSciences grant DE-FG02-92ER14244.

