## Supplemental figures for "Synaptic Zn^2+^ potentiates the effects of cocaine on striatal dopamine neurotransmission and behavior"

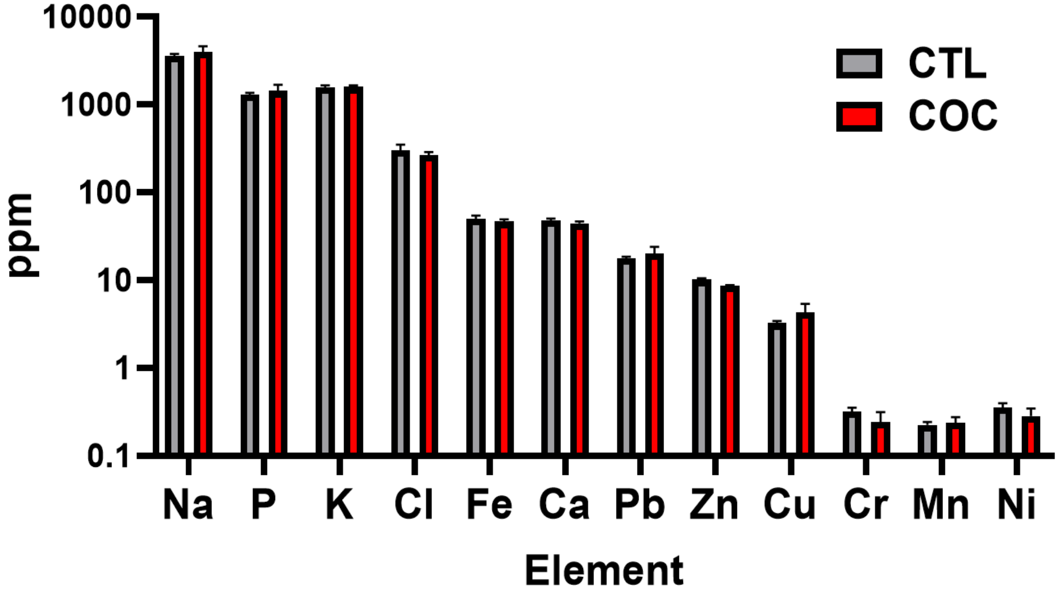


**Fig. S1. Elemental profiling in postmortem striatal tissue from cocaine users (COC) (n=19) and control (CTL) (n=20) subjects using total reflection X-ray fluorescence spectroscopy. Ppm – parts per million. Data expressed as Mean ±SEM.**

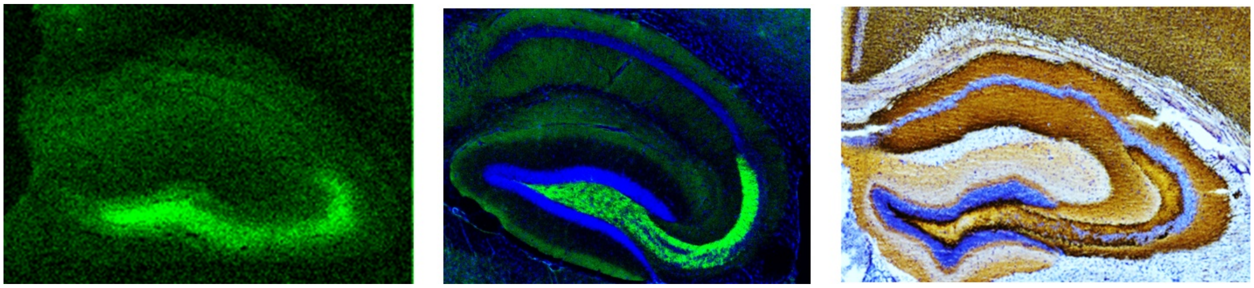

**Fig. S2. Visualization of synaptic $Zn^{2+}$ using synchrotron X-ray fluorescence microspectroscopy (XRFS). XRFS (left) signal overlaps with ZnT3 immunohistochemistry (middle), and histochemically-reactive synaptic $Zn^{2+}$ stain (right) in hippocampus.**

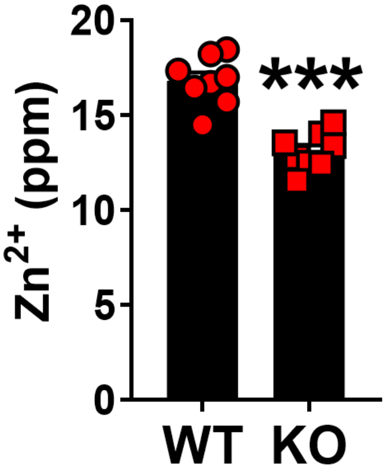


**Fig. S3. ZnT3 knockout (KO) mice have significantly lower (unpaired t-test; $t=6.419$; $p<0.001$) $Zn^{2+}$ in cortex compared to wildtype (WT) mice as assessed using total reflection X-ray spectroscopy (TXRF). ***$p \leq 0.001$. Data expressed as Mean ±SEM.**

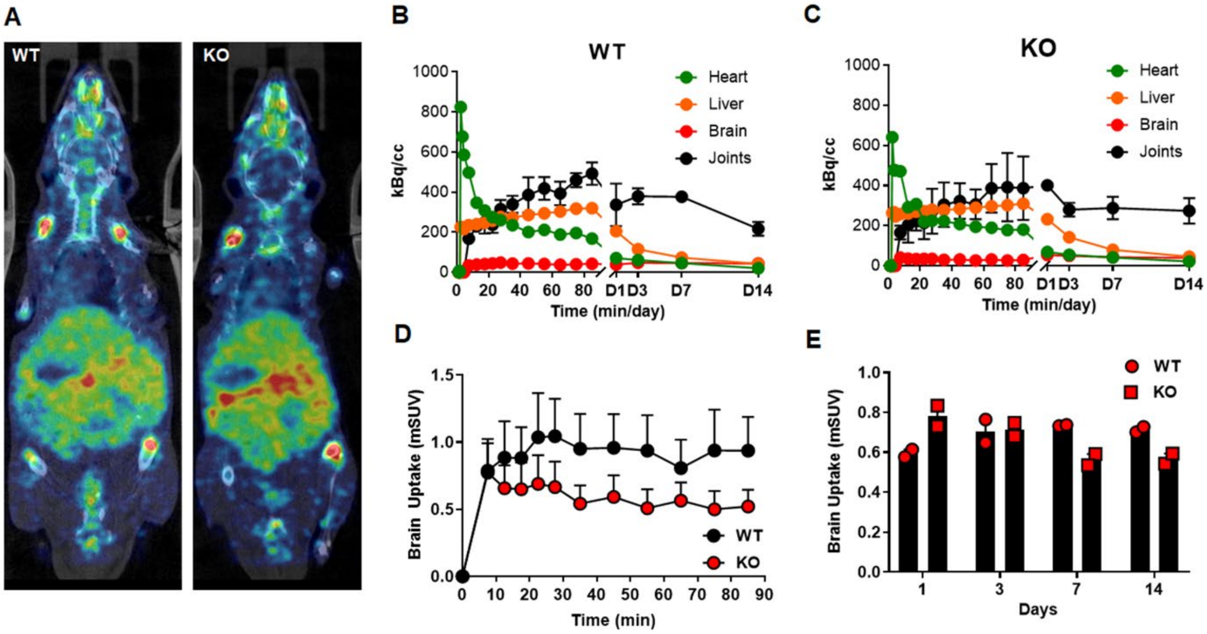


**Fig. S4.** **(A) Representative whole-body PET images (horizontal plane) of $^{65}$Zn uptake in a wildtype (WT) and a ZnT3 knockout (KO) mouse. (B) $^{65}$Zn time activity curves in different organs or body regions in WT and (C) ZnT4 KO mice. (D) $^{65}$Zn brain uptake over the first 90 min after $^{65}ZnCl_2$ intravenous injection. (E) $^{65}$Zn brain uptake at different days after $^{65}ZnCl_2$ intravenous injection showing that KO mice differed in brain uptake at 1, 7, and 14 days. Data expressed as Mean ±SEM.**

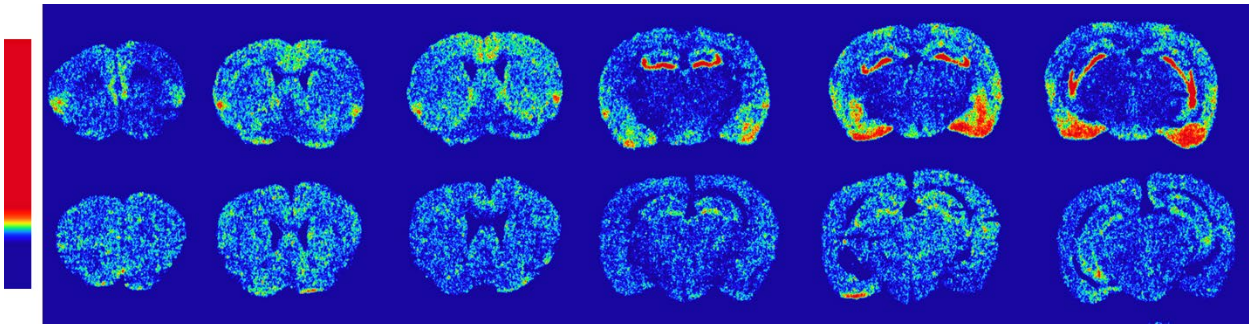

**Fig. S5.** ***Ex vivo*** **autoradiography at 15 days after intravenous $^{65}$Zn injection in a wildtype (top row) and a ZnT3 knockout mouse (bottom row).**

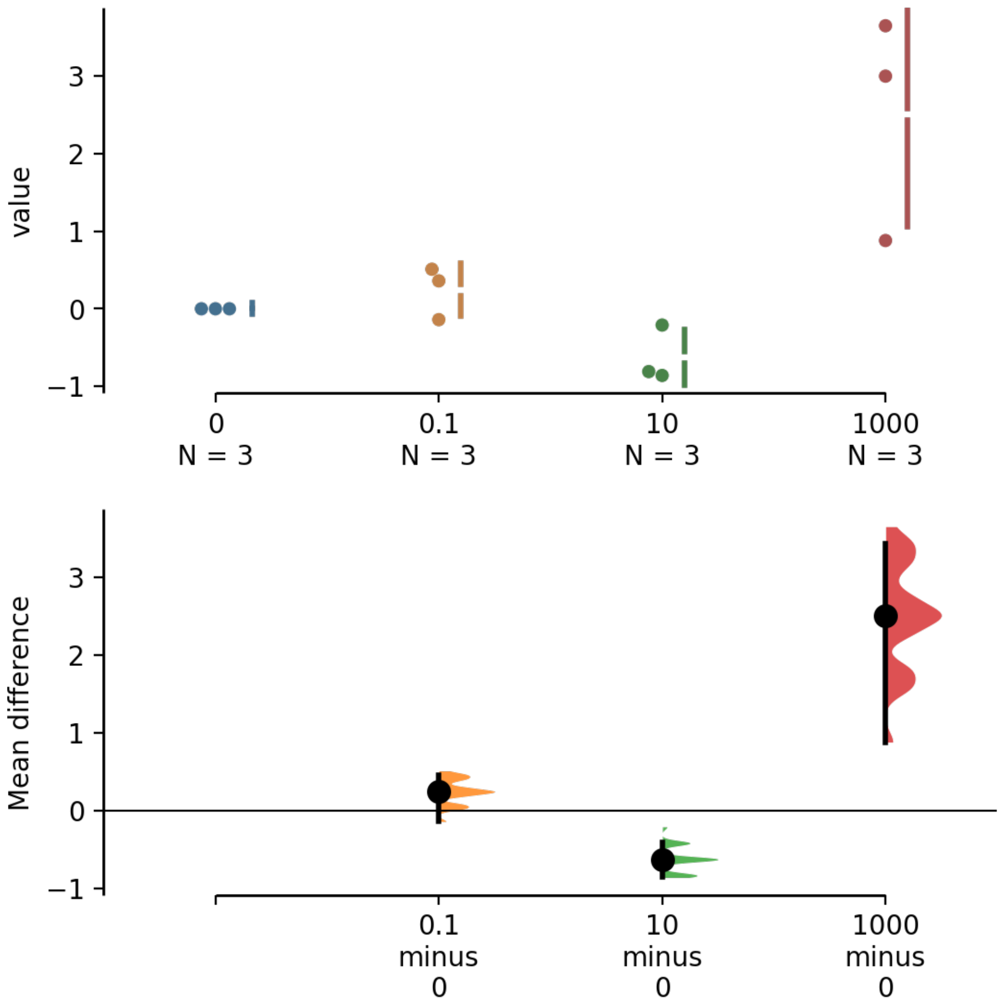


**Fig. S6. Cumming estimation plot showing the mean difference for the 3 comparisons of $Zn^{2+}$ concentrations and their effects on [$^3$H]WIN35,428 binding affinity (0 vs. 0.1 µM, 0 vs. 10 µM, 0 vs 1000 µM).**

| | WT | KO | ANOVA |
|---|---|---|---|
| **$DA_{Max}$** | 0.042 ± 0.007 µM (n=4) | 0.098 ± 0.035 µM (n=4) | $F_{1,6}$ = 2.4, p = 0.172 |
| **Clearance (k)** | 2.8 ± 0.6 µM/s (n=4) | 1.5 ± 0.4 µM/s (n=4) | $F_{1,6}$ = 3.8, p = 0.099 |

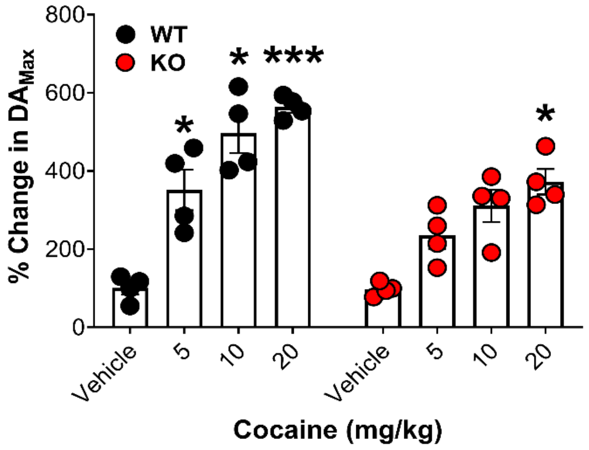


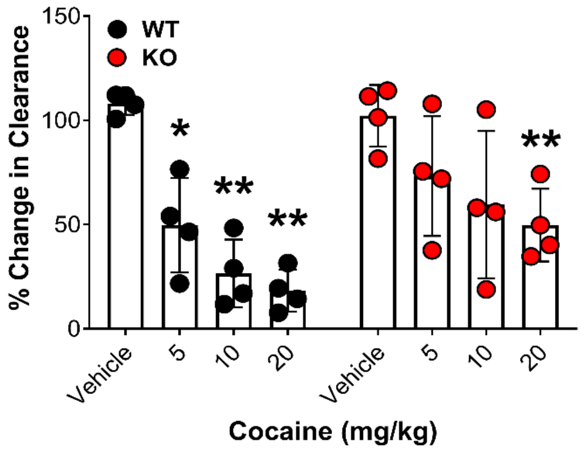


**Fig. S7. <u>Upper:</u> Table showing baseline fast scan cyclic voltammetry measures and statistics of $DA_{MAX}$ and Clearance (k) in wild-type (WT) and ZnT3 knockout (KO) mice. <u>Lower:</u> Data from each 5 min bin from Figure 3 were averaged per treatment (e.g., vehicle, 5, 10 and 20 mg/kg). *$DA_{MAX}$:* A 2-way repeated measures (RM) ANOVA (significant interaction effect: F(3, 18)=6.42; p=0.003) showed that WT mice had significantly greater $DA_{MAX}$ (Holm-Sidak multiple comparisons) at 5 mg/kg (t=5.39; p=0.03), 10 mg/kg (t=8.18; p=0.01) and 20 mg/kg (t=31.1; p<0.001) cocaine as compared to Vehicle, whereas ZnT3 KO mice had significantly greater $DA_{MAX}$ (Holm-Sidak multiple comparisons) only at 20 mg/kg (t=7.63; p=0.02) cocaine as compared to Vehicle. *Clearance:* A 2-way RM ANOVA (significant interaction effect: F(3, 18)=5.44; p=0.007) showed that WT mice had significantly greater Clearance (Holm-Sidak multiple comparisons) at 5 mg/kg (t=6.53; p=0.02), 10 mg/kg (t=10.75; p=0.008) and 20 mg/kg (t=17.44; p=0.002) cocaine as compared to Vehicle, whereas ZnT3 KO mice had significantly greater Clearance (Holm-Sidak multiple comparisons) only at 20 mg/kg (t=9.73; p=0.01) cocaine as compared to Vehicle. *p<0.05, **p<0.01, ***p<0.001. All data shown as Mean ±SEM.**

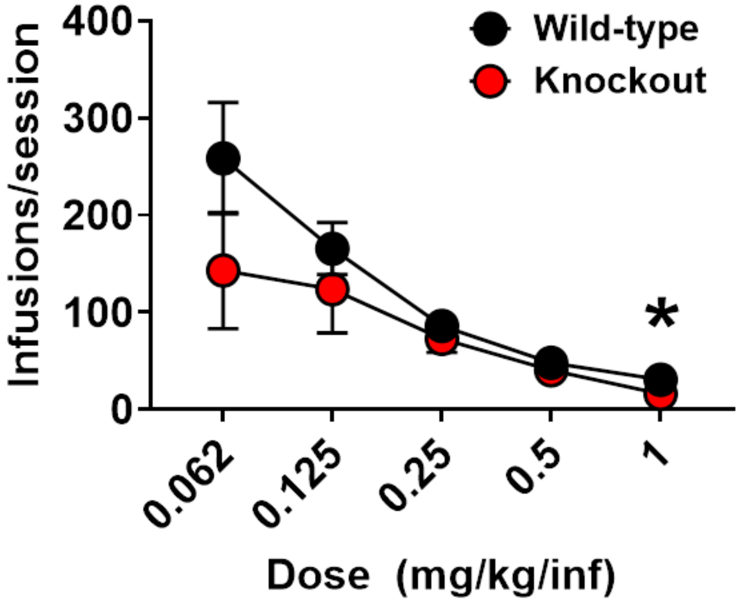


**Fig. S8. Infusions per session across different doses of intravenous cocaine self-administration between wild-type and ZnT3 knockout mice (t=2.54; p=0.03 at 1 mg/kg/infusion). Data expressed as Mean ±SEM.**
